# Müllerian mimicry in Neotropical butterflies: One mimicry ring to bring them all, and in the jungle bind them

**DOI:** 10.1101/2025.01.30.635679

**Authors:** Eddie Pérochon, Neil Rosser, Krzysztof Kozak, W. Owen McMillan, Blanca Huertas, James Mallet, Jonathan Ready, Keith Willmott, Marianne Elias, Maël Doré

**Affiliations:** Institut de Systématique, Evolution, Biodiversité, MNHN-CNRS-Sorbonne Université-EPHE-Université des Antilles, Muséum national d’Histoire naturelle de Paris, 45 Rue Buffon, 75005, Paris, France; Department of Ecology and Evolution, University of Lausanne, Lausanne, Switzerland; Department of Biology, University of Miami, Coral Gables, Florida 33146, USA; Museum of Comparative Zoology, Harvard University, Cambridge, Massachusetts 02138, USA; Smithsonian Tropical Research Institute, Panamá, Panamá; Museum of Vertebrate Zoology, UC Berkeley, USA; Department of Science, Natural History Museum, London, UK; Centre for Advanced Studies of Biodiversity, Instituto de Ciências Biológicas, Universidade Federal do Pará, Belém, Brazil; McGuire Center for Lepidoptera and Biodiversity, Florida Museum of Natural History, University of Florida, Gainesville, Florida, USA; Centre d’Ecologie et des Sciences de la Conservation, UMR 7204 MNHN-CNRS-Sorbonne Université, Muséum national d’Histoire naturelle de Paris, 45 rue Buffon, 75005, Paris, France

**Keywords:** biodiversity hotspots, comparative phylogenetic analyses, heliconiines, ithomiines, Müllerian mimicry, Neotropical butterflies, Neotropics, niche convergence, spatial co-occurrence

## Abstract

Understanding the mechanisms underlying species distributions and coexistence is essential to predict and prevent the impacts of global change, particularly in biodiversity hotspots. However, the effects of biotic interactions may be challenging to investigate at large spatial scales. Leveraging well-characterized Müllerian mimetic systems in Neotropical butterflies, we investigated spatial patterns of mutualistic mimetic interactions within and between two tribes of aposematic Nymphalid butterflies: the Heliconiini (Heliconiinae) and the Ithomiini (Danainae). Despite 85 My of independent evolutionary histories, many species share similar warning wing patterns across the Neotropics.

In this study we show that both tribes form similar biodiversity hotspots with a high prevalence of rare species and mimetic patterns in the tropical Andes. However, we reveal a higher relative richness of heliconiine butterflies than ithomiines in the Amazon basin contrasting with the Andean concentration of ithomiine diversity. Despite this difference in broadscale diversity patterns, we also document large-scale spatial associations among phenotypically similar species within and between tribes, thereby providing new empirical evidence for Fritz Müller’s historical model of mutualistic mimicry at a continental scale. Furthermore, comparative phylogenetic analyses suggest that co-mimetic species within and between tribes have converged towards similar climatic niches as a response to selection favoring co-occurrence.

Our findings illustrate the strength of mutualistic interactions in shaping biodiversity patterns at continental scale and in supporting niche convergence even across millions of years of evolution. Critically, it also emphasizes the pervasive vulnerability of mimetic communities, bound by positive interactions, to disassembly induced by climate change.

**Significance statement:** Müllerian mimicry is a remarkable example of convergent evolution driven by natural selection where coexisting prey species converge in their warning signal advertising their defenses to predators. Heliconiine and ithomiine butterflies found throughout Neotropical rainforests were instrumental in Fritz Müller’s historical model, which provided the mechanism for such resemblance. Leveraging decades of fieldwork and museum collections, we show that species with similar color patterns present strikingly similar spatial distributions, regardless of how closely related they are. Such co-occurrence appears reinforced by the evolution of similar climatic requirements among look-alike species. Our findings emphasize the key role of mutualistic interactions in shaping large-scale patterns of biodiversity and supporting convergence in the climatic niches of species spanning across phylogenetically distant clades.

## 1 Introduction

Biotic interactions are known to structure ecological communities (1) but their impact on biodiversity patterns remains difficult to quantify, especially at large spatial scales (2, 3). Biotic interactions include negative interactions such as exploitative competition for resources, positive interactions such as pollination, and asymmetrical interactions such as predation. Ecologists also distinguish intraguild interactions occurring between species in the same ecological guild, such as competition for similar resources, from interguild interactions between species in different ecological guilds, such as predators and prey. As such, interactions underlie numerous complex ecological and evolutionary processes and involve virtually all life forms (1).

Extensive theoretical and empirical evidence supports the role of negative intraguild interactions in driving spatial and phenotypical divergence among competing species (4–7). By contrast, intraguild mutualistic interactions remain some of the most understudied, yet they can also have important consequences for both trait evolution and the geographic distributions of species involved (8). For instance, selection may favor evolutionary convergence in flowering phenology as well as floral traits that allow different plant species to benefit from attracting a similar group of pollinators (9–12). Furthermore, facilitation shapes the distribution of plant species allowing the co-occurrence of distantly related species, thus enhancing phylogenetic diversity (13, 14). However, such potentially broadscale effects of intraguild mutualistic interactions remain largely overlooked outside of plants and microorganisms (15, 16).

In this study, we investigate the consequences of Müllerian mimicry for species niche evolution and community composition at large spatial and phylogenetic scales. Müllerian mimicry occurs between coexisting prey species that have evolved similar warning aposematic patterns advertising their defenses to predators (17, 18). Such intraguild interactions are mutualistic because species displaying the same warning pattern benefit from sharing the mortality cost of educating naive predators among a larger set of individual prey (19). Müllerian mimicry has been described in many organisms, including birds, insects, snakes, fishes, and amphibians (20–26). Numerous independent origins across distantly related taxa reinforce the idea that Müllerian mimicry represents a selective advantage for defended prey (18, 19, 27). Unlike many ecological interactions, Müllerian mimicry is relatively straightforward to characterize: in a given community, defended prey species sharing a common warning signal form groups called ‘mimicry rings’ underlying mutualistic interactions (28–30), while species harboring different signals do not interact through mimicry.

Two diverse tribes of Neotropical nymphalid butterflies, the Heliconiini Swainson, 1822 and Ithomiini Godman & Salvin, 1879, provide an excellent study system to assess the effect of Müllerian mimicry on species distributions and ecological niches between two distantly related clades (**Fig. 1.b;** 31). Both tribes were instrumental in the discovery (32) and formalization of Müllerian mimicry (19), a pivotal finding that provided both empirical and theoretical supports for the then young theory of evolution by means of natural selection formulated concomitantly by Charles Darwin and Alfred Russel Wallace (33, 34). All species in Heliconiini and Ithomiini are considered to be chemically defended to varying degrees, thereby being unpalatable to predators (35–38). Heliconiine butterflies sequester toxic chemical compounds from their Passifloraceae host plants (39, 40) and/or can synthesize *de novo* compounds from amino-acids (41), whereas ithomiines mostly derive toxic compounds from feeding on decaying leaves or flowers of Boraginaceae and Asteraceae as adults (38, 42). Both tribes are widely distributed across the American continent, from Canada (Heliconiini) and Mexico (Ithomiini) to northern Argentina, and from the Pacific to the Atlantic and Caribbean coasts (43, 44). Throughout this wide range, many species interact both within and between the tribes via Müllerian mimicry.

**Figure 1.**
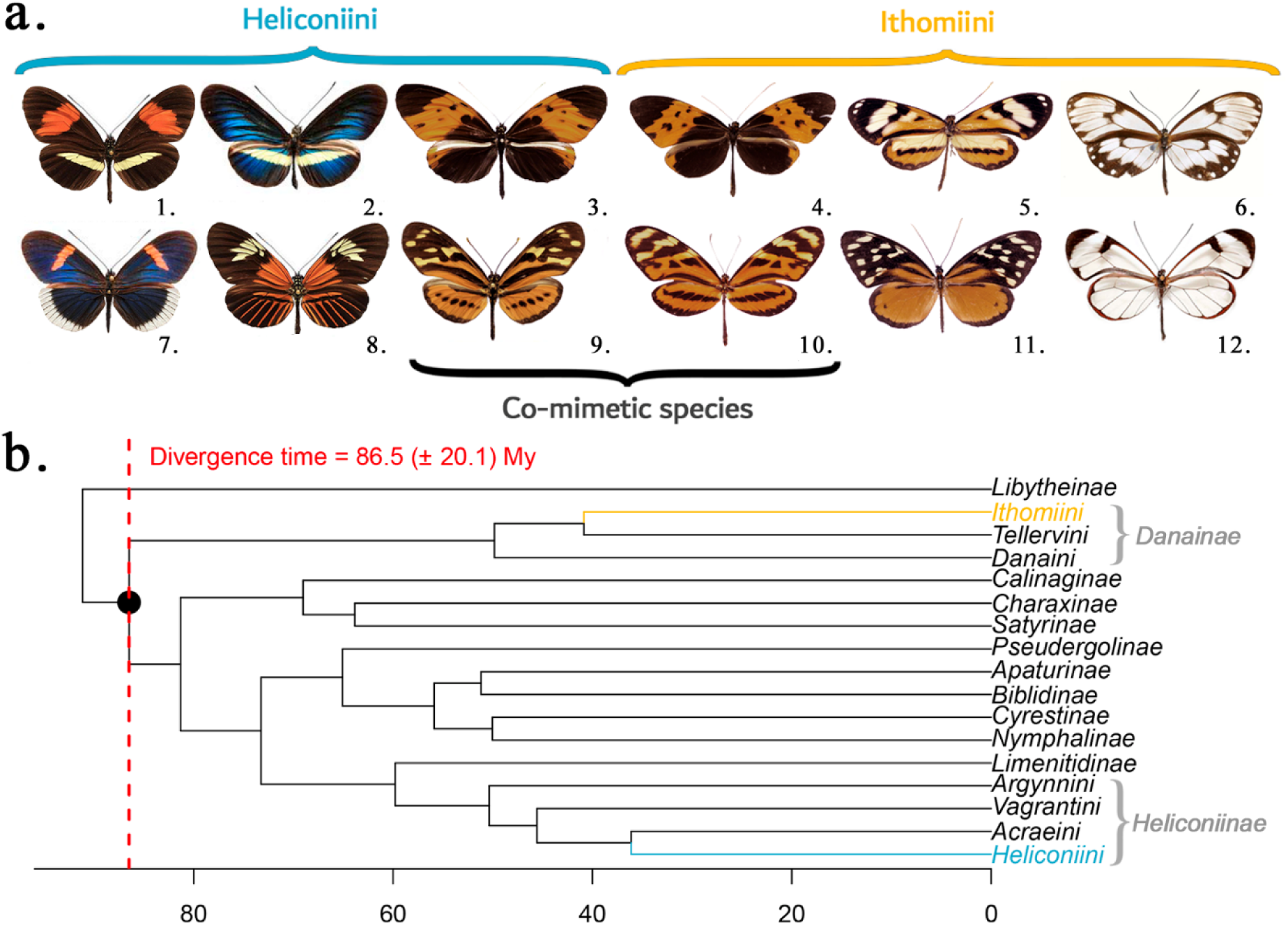
(a) Diversity of wing patterns within and between tribes in Ithomiini and Heliconiini. The four central butterflies represent examples of subspecies that share similar wing patterns and take part in co-mimetic mutualistic interactions through Müllerian mimicry. From 1 to 12: *Heliconius melpomene amaryllis*, *Heliconius erato chestertonii*, *Heliconius numata bicoloratus, Melinaea isocomma simulator*, *Hypothyris ninonia daeta*, *Veladyris pardalis christina*, *Heliconius sapho sapho*, *Heliconius elevatus elevatus*, *Eueides isabella dissoluta*, *Mechanitis lysimnia utemaia, Tithorea harmonia helicaon*, *Greta morgane oto*. Comprehensive plates of the 38 heliconiine phenotypic groups (**Fig. S1**) and 44 ithomiine phenotypic groups (**Fig. S2**) are available in **Appendix 1**. **(b) Relative position of Ithomiini and Heliconiini tribes in the Nymphalidae phylogeny.** Extracted from Chazot et al. (31). Tip labels represent butterfly subfamilies except for Danainae and Heliconiinae, which are divided into tribes. The red-dotted line represents the estimated divergence time between Heliconiini and Ithomiini.

The tribe Heliconiini contains 8 genera, ca. 77 species and 457 subspecies (45, 46, but see Núñez et al. (47) for recent proposed taxonomic updates). The tribe Ithomiini contains 42 genera, 396 species and 1542 subspecies (44, 48), with many having partially transparent wings, such as the emblematic Glasswing butterfly *Greta morgane oto* (**Fig. 1.a**). Despite having diverged about 86 million years ago (**Fig. 1.b;** 31), about the same time that humans split from flying lemurs (Order: Dermoptera; 49), the two tribes share numerous warning patterns and thus interact through mimicry (**Fig. 1.a**).

Recent work showed that mutualistic interactions have led to large-scale spatial associations and climatic niche convergence between phenotypically similar species in ithomiine butterflies (50). Here, we extend that scope to investigate the effects of mimicry between the distantly related tribes Heliconiini and Ithomiini on their biodiversity patterns at continental scale. While biogeographic patterns of species richness in ithomiine and heliconiine butterflies are already known (43, 44), no study has yet investigated the multiple facets of biodiversity patterns of both tribes in an integrated framework. Such an integrative approach encompassing the two most diverse adaptive radiations of Neotropical mimetic butterflies (44, is crucial to jointly define mimicry rings in local butterfly communities and provides statistical support for the co-occurrence of phenotypically similar butterfly species throughout the Neotropics. From an evolutionary perspective, it allows a better understanding of the role of mimetic interactions in shaping large-scale distribution patterns, and niche and trait evolution across phylogenetically distant clades (51).

Specifically, we aim to:

(1) Predict heliconiine subspecies distributions, map broadscale biodiversity patterns of Heliconiini including species richness, phylogenetic diversity, geographic rarity, and phenotypic richness, and test for congruences and disparities with known Ithomiini biodiversity patterns.
(2) Test whether phenotypically similar species co-occur at large spatial scales, supporting pervasive mutualistic interactions between Heliconiini and Ithomiini.
(3) Test if mutualistic interactions are associated with the convergence of the climatic niche of phenotypically similar species within and between tribes throughout the Neotropics.

## 2 Results

We classified heliconiine subspecies into 38 phenotypic groups based on wing pattern similarity (**Fig. S1** in **Appendix 1**). Because those groups are defined only on phenotypic similarity, members of such groups may not currently be involved in mutualistic interactions as they may not actually co-occur. Each phenotypic group therefore represents ‘putative’ local mimicry rings, as in Doré et al. (50). If a significant signal of spatial co-occurrence within a phenotypic group is detected, it then qualifies as an ‘effective mimicry ring’ depicting current ecological interactions (52, 53).

### 2.1 Congruence and contrasts in biodiversity patterns between tribes

While we found significant correlations in global biodiversity patterns and location of hotspots, we also detected notable regional differences between the two tribes (**Tables S1** in **Appendix 4**; see **Fig. S3** in **Appendix 2** for a map of bioregions). Similar to Rosser et al. (43), we found a peak of species richness in the eastern slopes of the Andes, with up to nearly half of the species in the group predicted to be found in some 30 km × 30 km grid cells (**Fig. 2.a**; 35 out of 77 species = 45.5%; **Fig. S4.a** in **Appendix 3**). We also detected secondary hotspots of species richness in the Amazon basin and in southern Central America. These broad-scale richness patterns are significantly correlated with those of Ithomiini (Spearman’s rho = 0.771, t-stat = 7.91, Clifford’s df = 42.7, Q95% = 1.681, p < 0.001; **Table S1** in **Appendix 4**), which also shows a peak of species richness in the Andes and southern Central America (44). However, Ithomiini have proportionally fewer species in the Amazon basin and the Brazilian Atlantic Forest (**Fig. 2.a)**. Finally, Heliconiini are present in the Nearctic region, while Ithomiini are only barely found north of Mexico (**Figs. S4.a & S5.a** in **Appendix 3**).

**Figure 2:**
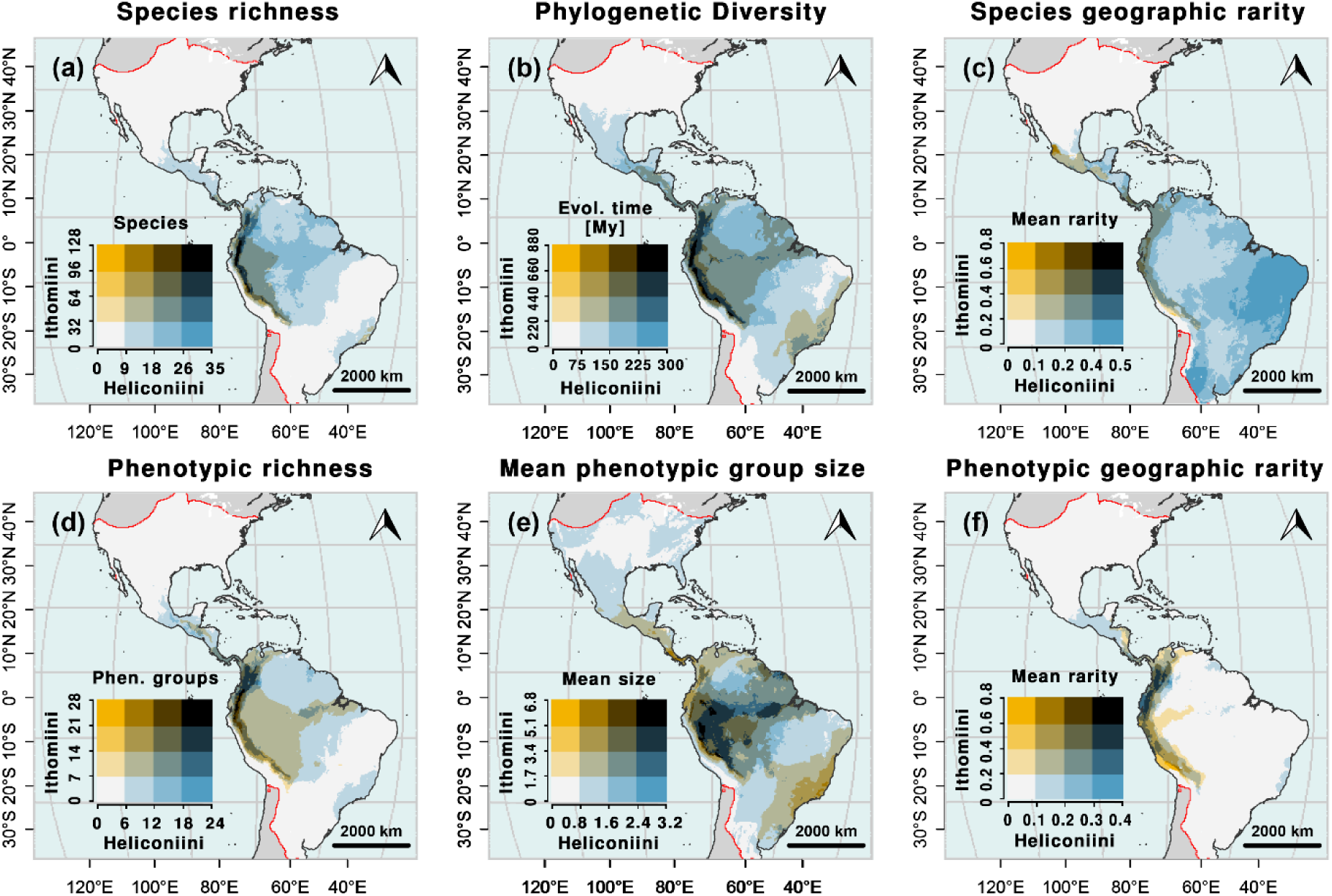
Relative patterns of biodiversity in Heliconiini and Ithomiini at the continental scale. **(a)** Species richness. **(b)** Faith’s Phylogenetic Diversity (55) **(c)** Mean species geographic rarity. Rarity index based on species ranges. **(d)** Phenotypic richness (i.e., number of local phenotypic groups). **(e)** Mean phenotypic group size (i.e., mean number of species per local phenotypic group). **(f)** Mean phenotypic geographic rarity. Rarity index based on phenotypic group ranges. The bivariate color scale represents the scaled values of each index in the two tribes. Values are scaled by the maximum value for each tribe; thus, they describe relative patterns of biodiversity. Blue areas reflect higher relative diversity/rarity for Ithomiini while yellow areas reflect higher relative diversity/rarity for Ithomiini. Darker areas represent shared hotspots of diversity/rarity. Ithomiini patterns are extracted from Doré et al. (44). Absolute patterns of Heliconiini and Ithomiini biodiversity can be found in **Figs. S4 & S5** in **Appendix 3.**

Phylogenetic diversity in Heliconiini was highly correlated with species richness (Spearman’s rho = 0.987, t-stat = 24.2, Clifford’s df = 15.5, Q95% = 1.749, p < 0.001; **Table S2** in **Appendix 4**), with the highest values found in the Andes and around the Amazon River, and with secondary hotspots in Central America and the Brazilian Atlantic Forest (**Fig. 2.b; Fig. S4.b** in **Appendix 3**). However, while Ithomiini phylogenetic diversity was mostly concentrated in the Andes (**Fig. 2.b**; **Fig. S5.b** in **Appendix 3**; 44), less spatial variance was predicted for Heliconiini (Coefficients of Variation asymptotic test: CV_Heliconiini_ = 0.395, CV_Ithomiini_ = 0.633, χ² = 3.98, Clifford’s sample size = 92.5, df = 1, Q95% = 3.841, p = 0.046), with values across the Amazon basin approaching those in the Andean regions (**Fig. S4.b** in **Appendix 3)**.

The Caatinga, a habitat with semi-arid tropical vegetation in Northeast Brazil, hosts the highest proportion of heliconiine species with restricted geographical ranges (**Fig. 2.c; Fig. S4.c** in **Appendix 3**). Though, this pattern is produced by only a couple of taxa that are restricted to this region (54). Meanwhile, geographical rarity was lowest in the Nearctic, which also hosts only a few species, but with wide geographical distributions, such as *Dryas iulia* and *Agraulis vanillae* (but see Núñez et al. (47) for recent proposed taxonomic splits). By contrast, the highest proportion of Ithomiini species with restricted ranges are found in the Andes and in Central America (**Fig. 2.c; Fig. S5.c** in **Appendix 3**; 44), leading to a lack of significant correlation in broadscale patterns of geographic rarity between the two tribes (Spearman’s rho = -0.042, t-stat = -0.34, Clifford’s df = 64.7, Q95% = 1.669, p = 0.632; **Table S1** in **Appendix 4**).

Spatial patterns of phenotypic richness in heliconiines (i.e., the number of ‘phenotypic groups’ represented at a given location) were strongly correlated to those of heliconiine species richness (Spearman’s rho = 0.978, t-stat = 18.0, Clifford’s df = 14.7, Q95% = 1.755, p < 0.001; **Table S2** in **Appendix 4**) and phenotypic richness in ithomiines (Spearman’s rho = 0.762, t-stat = 7.60, Clifford’s df = 41.8, Q95% = 1.682, p < 0.001; **Table S1** in **Appendix 4**). Both tribes displayed a sharp contrast between the Andes and the rest of the continent. Maximum phenotypic richness was reached in the northern Andes, where up to 24 (63.2%) of the 38 Heliconiini color patterns and 29 (65.9%) of the 44 Ithomiini patterns are predicted to be found in local grid cells (**Fig. 2.d; Fig. S5.d** in **Appendix 3**; 44). Phenotypic richness in the Western Amazon basin stands out much less for Heliconiini than for Ithomiini (**Fig. 2.d; Fig. S4.d** in **Appendix 3**), because of a higher number of local species sharing similar color patterns compared to the Andes. Phenotypic groups of Heliconiini in the Amazon comprise between 1.5 to 3 species on average, but only up to 1.5 species in the Andes (**Fig. 2.e; Fig. S4.e** in **Appendix 3**). Furthermore, high phenotypic geographic rarity reflects the presence of phenotypic groups with smaller distribution ranges in the Andes compared to the rest of the continent (**Fig. 2.f**). Meanwhile, ithomiines form larger phenotypic groups in the Andes, the western Amazon, Central America, and the Brazilian Atlantic Forest, with between 3.5 to 7 species per group on average (**Fig. 2.e; Fig. S5.e** in **Appendix 3**; 44). However, ithomiine and heliconiine phenotypic groups are fairly similar to each other in terms of geographic rarity of mimicry patterns (Spearman’s rho = 0.625, t-stat = 5.71, Clifford’s df = 50.9, Q95% = 1.675, p < 0.001; **Table S1** in **Appendix 4**), with widely distributed patterns in Amazonia and patterns with narrow distributions in the Andes.

### 2.2 Mimicry promotes broadscale spatial congruence of phenotypically similar species

To explore whether mutualistic interactions can shape the co-occurrence of phenotypically similar species, within and between tribes, we used the Bray-Curtis (BC) index to quantify dissimilarities in large-scale spatial patterns of species. We compared the observed mean BC within phenotypic groups against BC obtained from random permutation of patterns between species as a null hypothesis depicting the absence of relationship between color patterns and spatial distributions of species. We detected that Heliconiini (Permutation test: BC_obs_ = 0.717, BC_null_ Q5% = 0.872, p ≤ 0.001), Ithomiini (Permutation test: BC_obs_ = 0.896, BC_null_ 5% = 0.946, p ≤ 0.001; 50) and inter-tribe phenotypic groups (Permutation test: BC_obs_ = 0.875, BC_null_ Q5% = 0.930, p ≤ 0.001) all display significantly lower mean spatial dissimilarities than at random (**Fig. S6** in **SI Appendix 5)**. As such, we detected a significant congruence of large-scale spatial distributions among phenotypically similar species, within and between tribes.

Tests were also carried out for each phenotypic group with at least two species. We observed that 14 out of 29 (48.3%) groups displayed significant signal for spatial congruence within the Heliconiini, supporting their qualification as ‘effective mimicry rings’ representing current mutualistic interactions. This proportion rises to 32 out of 39 (82.1%) groups within the Ithomiini (50). For color patterns shared between the two tribes, 9 out of 11 (81.8%) showed significant spatial congruence and are thus supported as ‘effective mimicry rings’ (**Table S3** in **Appendix 6**). For instance, the pattern EXCELSA showed an important and significant overlap of distributions between tribes throughout Central America and the Northern Andes (**Fig. 3.a**; Permutation test: BC_obs_ = 0.805, BC_null_ Q5% = 0.894, p ≤ 001),). Similarly, the PAVONII pattern appeared confined to the Northern and Central Andes for both tribes (**Fig. 3.b**; Permutation test: BC_obs_ = 0.653, BC_null_ Q5% = 0.819, p ≤ 001), while the heliconiines and ithomiines harboring the pattern MAELUS showed significant overlap in distributions across the Amazon Basin (**Fig. 3.c**; Permutation test: BC_obs_ = 0.643, BC_null_ Q5% = 0.894, p ≤ 001).

**Figure 3:**
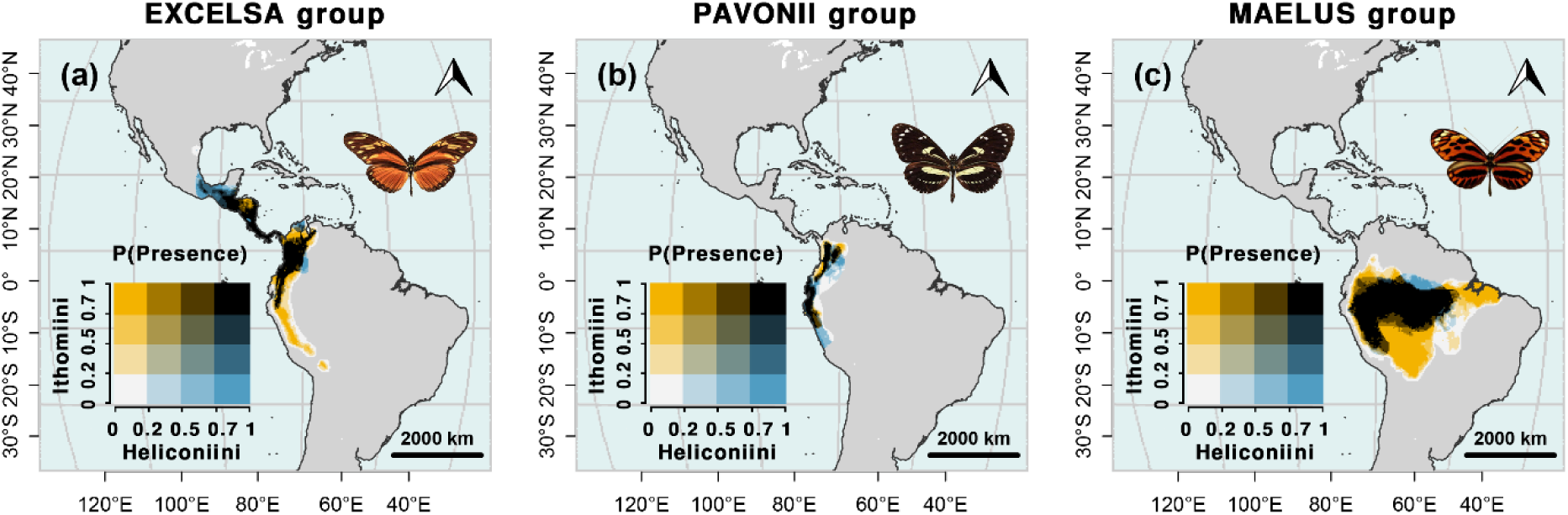
Comparative distributions of predicted presence for Heliconiini and Ithomiini phenotypic groups defined based on phenotypic similarity. **(a)** EXCELSA group. **(b)** PAVONII group. **(c)** MAELUS group.

### 2.3 Mimicry is associated with climatic niche convergence of phenotypically similar species

Beyond spatial distributions, we investigated the extent and significance of climatic niche convergence between phenotypically similar species for each type of phenotypic group (Heliconiini, Ithomiini, inter-tribe groups). We quantified climatic distances between species as Euclidean distances between the centroids of species occurrences in the bioclimatic space employed for Species Distribution Modeling (SDM). We compared the observed mean climatic niche distance (i.e., MCD) within phenotypic groups against MCD obtained from simulations of neutral evolution of the climatic niche along the phylogeny. The observed MCD was significantly lower for all three types of phenotypic groups (Heliconiini only: MCD_obs_ = 0.684, MCD_null_ Q5% = 0.882, p ≤ 001; Ithomiini only: MCD_obs_ = 0.724, MCD_null_ Q5% = 0.954, p ≤ 001 adapted from Doré et al. (50); inter-tribe: MCD_obs_ = 0.725, MCD_null_ Q5% = 0.866, p ≤ 0.001). Therefore, we detected a significant evolutionary association between climatic niche and color patterns, both within and between tribes, as species sharing wing patterns tend to have more similar climatic niches than expected under neutral niche evolution.

Overall, we observed that 12 out of 29 (41.4%) phenotypic groups in Heliconiini, and 33 out of 39 (84.6%) phenotypic groups in Ithomiini (50), showed significant niche convergence. For inter-tribe phenotypic groups, niche convergence was supported for 6 out of 10 (60.0%) groups (**Table S4** in **Appendix 6**).

## 3 Discussion

Our study highlights the strength of mutualistic interactions to shape biodiversity patterns at continental scale and to support species niche convergence across evolutionarily distant lineages. Specifically, we showed how Müllerian mimicry shapes large-scale spatial congruence in distributions and niche convergence within and between two emblematic tribes of unpalatable Neotropical butterflies that diverged from a common ancestor roughly 85 million years ago.

Both heliconiine and ithomiine butterflies display a high diversity and concentration of species and color patterns in the tropical Andes, with an important proportion displaying geographically restricted ranges. Although we detected minor differences in regional biodiversity patterns between the two tribes, likely because of differences in host plant distributions and biogeographic origins (see below), we provided further evidence that mimicry is associated with large scale spatial congruences among phenotypically similar species, both within and between tribes. This finding provides new empirical evidence for the unfolding of the prediction of Müller’s mimicry model at a macroecological scale and across millions of years of evolution. Furthermore, comparative phylogenetic analyses suggest that mimetic interactions support the evolutionary association between climatic niche and color pattern evolution, within and even across tribes, as a consequence of selection favoring both the phenotypic convergence of sympatric species and the co-occurrence of look-alike species.

### 3.1 Drivers of global biodiversity patterns

Our analyses predict species richness and phylogenetic diversity of Heliconiini to be particularly high in the Amazon basin and in the tropical Andes (**Fig. 2**), in line with previous findings (43, 56). Ithomiini also exhibit high species and phylogenetic diversity patterns in these regions, especially in the tropical Andes (44, 57; **Fig. S5** in **Appendix 3**). Those congruent biodiversity patterns are similar to those observed in other taxa, including angiosperms (58), beetles (59), birds (60), mammals (61), reptiles (62), and amphibians (63), reinforcing the position of the tropical Andes as the richest biodiversity hotspot on Earth (63, 64). This outstanding biodiversity is strongly influenced by geological and climatic factors. The topographical complexity of the tropical Andes facilitates fine-scale spatial variation of environmental conditions and provides more opportunities for parapatric and allopatric speciation, fueling regional adaptive radiations (65–67). Moreover, the tropical Andes and the Amazon basin have benefited from a historically relatively stable climate, thought to reduce species extinction rates (68, 69) and allow for the long-term persistence of high levels of species diversity and endemism (70–73). Noticeably, the Brazilian Atlantic Forest (Mata Atlântica), although geographically distant from the Andes and the Amazon basin, also harbors a relatively high diversity of both Heliconiini and Ithomiini. This disjunct pattern could be the result of multiple independent colonization events resulting from changing connectivity with Amazonia (48, 74, 75), followed by local diversification in the Brazilian Atlantic Forest.

While broadscale diversity patterns of Heliconiini are significantly correlated with those of Ithomiini (**Table S1** in **Appendix 4**), we also observed minor regional disparities between the two tribes. Ithomiini richness and rarity is concentrated in the tropical Andes (44, 57), while heliconiines harbor high richness in both the Andes and the Amazon basin (**Fig. 2.a**). This difference could be explained by contrasted geographic origins between the tribes. Ithomiini are predicted to have originated in the Central Andes (48), allowing time for speciation and lineage accumulation in this region. Formal historical biogeographic reconstructions at the tribe level are still lacking for Heliconiini (76), but an Amazonian origin of Heliconiini could explain their relative richness in the area. Perhaps more importantly, both tribes encompass numerous species that typically feed on a small number of larval host plants (77, 78). However, they specialize on distinct families of plants, the Passifloraceae for the heliconiines and the Solanaceae for most ithomiines, with which they are suspected to have tightly coevolved (46, 77). Thus, differences in biogeographic histories and species richness patterns of the two host plant lineages (79–81) could partly explain the current dissimilarities observed in biodiversity patterns between the two butterfly tribes.

Beyond taxonomic and phylogenetic patterns, both tribes exhibit contrasting mimetic characteristics between Andean and Amazonian communities. A few large-range and species-rich phenotypic groups dominate mimetic butterfly communities in the Amazon basin (**Fig. 2.d-f**). This suggests a pervasive wing pattern convergence between distant lineages and/or high degree of phenotypic conservatism within local radiations acting alongside relatively strong frequency-dependent selection purging any less common patterns that may arise across the relatively homogeneous climate of the Amazon basin (82). Conversely, in the Andes, where phenotypic groups have smaller distributions and are composed of fewer species, there is a higher diversity of wing patterns per grid cell (**Fig. 2.d-f**). This is likely explained by strong environmental gradients and geographic barriers present in these mountainous regions. These abiotic features favor partitioning of predator and prey communities across space, which incurs selection on color patterns through mimicry at a scale smaller than our 30 km × 30 km grid cells, thereby driving the partitioning of species among more phenotypic groups (83).

### 3.2 Large-scale spatial congruence of phenotypically similar species

Despite showing some minor disparities in global biodiversity patterns, the two tribes are strongly linked through mimetic interactions. An important proportion of the phenotypic groups identified within each tribe, but also between the two tribes, show significant large-scale spatial congruence. Those groups likely represent ‘effective mimicry rings’ as sets of phylogenetically distant but phenotypically similar species involved in mutualistic interactions in local communities. However, when studying the Heliconiini group PAVONII, we found no evidence of co-occurrence nor climatic niche convergence among species for this phenotypic group. Only after accounting for the Ithomiini members of this inter-tribe phenotypic group did we detect an overall significant signal for spatial congruence (**Fig. 3.b**; **Tables S3 & S4** in **Appendix 6**). Therefore, our study highlights the importance of accounting for distinct members of the mimetic community when investigating how mutualistic interactions shape the distribution of species, even when those are distantly related.

### 3.3 Climatic niche associations over 85 My of independent evolution

Beyond similarities in spatial distributions, we found a significant evolutionary association between phenotypic patterns and species climatic niche both within and between tribes. Patterns of trait and niche convergence across co-mimetic species of Neotropical butterflies have already been detected for flight behavior (51), and for ecological dimensions acting at local scales, such as nocturnal roosting habitat height (84), flight height and microhabitat (85–88), and forest structure (88, 89). Here we show that convergence can arise for niche dimensions (i.e., climatic niche) that directly affect biodiversity patterns at the continental scale and can link the fate of two tribes that, despite being separated by over 85 My of independent evolution, currently share highly similar phenotypes, spatial distributions, and associated climatic niches.

Such strong adaptive bonding across tens of millions of years of evolution may have significant implications in the face of the ongoing climate change. Indeed, Müllerian mimicry represents interactions that are beneficial for the individuals involved, compensating for the negative effects of resource and habitat competition (90), and fueling higher local species richness (91, 92). However, if mutualistic interactions are lost because of species extinction or community disassembly, their disappearance can reduce community stability and potentially trigger cascades of local extinction (93, 94). The dispersal abilities of Müllerian mimetic species are impeded by the purifying selection acting on individuals harboring novel phenotypes in newly colonized areas (27, 95). Moreover, despite relatively similar climatic niche optima, tolerance to climate fluctuations and extremes and species dispersal abilities may still differ among co-mimetic species, limiting opportunities for co-dispersal trajectories and leading to community disassembly (71, 96, 97). Finally, the effects of climate change on biotic factors that affect local abundance, such as host plants (98, 99) and parasitoids (100, 101), may also differ among interacting species, impeding even more their abilities to cope with climate change as tightly coevolved assemblages are tied together by positive interactions (102).

### 3.4 Limitations and perspectives

We found a statistical support for an evolutionary association of species climatic niche and aposematic color patterns, within and between tribes. However, our analyses do not allow us to disentangle whether this pattern resulted from selection favoring the phenotypic convergence of sympatric species or the co-occurrence and associated niche convergence of look-alike species. In practice, both mechanisms are likely involved (50). Besides favoring the phenotypic convergence of sympatric species, the reduced cost of predation associated with Müllerian mimicry (19) may enable survival of rare forms within a species and assist the colonization of new environments shared with their mimetic partners (92), resulting in effective niche convergence. In the peculiar case of frequency-dependent Müllerian mimicry, such evolution likely occurs through advergence rather than gradual convergence, with the rarest species evolving towards the more abundant, thus better numerically defended, species (103). The most likely scenario in our system is advergence of heliconiines towards ithomiines, both because of color pattern ancestry, such as subspecies of *Heliconius numata* mimicking different species of *Melinaea* (104), and greater numerical abundances in ithomiines (32, 105).

While ithomiines and heliconiines form the bulk of butterfly mimetic communities in the Neotropics (105, 106), they also interact with a wide range of other mimetic butterflies and diurnal moths, such as the chemically defended Dioptinae (Notodontidae; 107) and Pericopina (Erebidae; 108), numerous presumed palatable Batesian mimics such as Dismorphiinae butterflies (Pieridae; 109), and even Polythoridae damselflies (105, 110, 111). How these relatively less explored, or still undiscovered, components of mimetic communities, and notably the existence of Müllerian and Batesian components, affect the whole distribution and niche evolution of interacting species, is virtually unknown (112, 113). Thus, future directions in this research topic may aim to enlarge the taxonomic scope and shed light on the importance of mimetic interactions in shaping large spatial biodiversity patterns across evolutionary distantly related lineages.

## 4 Materials and Methods

### 4.1 Phenotypic classification of wing color patterns

We classified heliconiine subspecies into 38 groups of wing color pattern similarity forming phenotypic groups (**Fig. S1** in **Appendix 1**) representing ‘putative’ local mimicry rings (as in Doré et al. (44, 50) for Ithomiini). Since those groups are formed based only on phenotypic similarity, members of such groups may not currently be involved in mutualistic interactions as they may not actually co-occur. If a significant signal of spatial co-occurrence within a phenotypic group is detected, it then qualifies as an ‘effective mimicry ring’, tentatively reflecting true ecological interactions (52, 53). We collected at least one digital image of the dorsal wing patterns for 436 out of 457 known subspecies of Heliconiini (Heliconiinae), taken from specimens held in museums and private collections. We clustered all Heliconiini images based on perceived similarity in their dorsal wing color, pattern, and shape. Geographic distributions of taxa were not considered during this process. For Ithomiini (Danainae), we used the currently accepted classification of mimicry patterns (44), built using a similar rationale of phenotypic similarity. Then, we matched the identity of phenotypic groups associated with a pattern represented in the two tribes and labeled them as inter-tribe phenotypic groups. In order to ensure the robustness of our results to alternative classifications, we also designed higher level groups encompassing multiple initial phenotypic groups. We carried out analyses for the two most extreme choices for the classification: the most ‘split’ with the 38 initial phenotypic groups as showed in the main text, and the most ‘lumped’ with 20 phenotypic groups (see **Appendix 7**). This design ensures that any intermediate choice in the phenotypic classification would lead to similar results, as long as results of the two extreme options lead to similar conclusions. The comprehensive phenotypic-based classification of heliconiine subspecies is available in an online archive (<u>10.5281/zenodo.10903197</u>).

### 4.2 Occurrence database and phylogenies

In order to map biodiversity patterns of heliconiine butterflies, we curated a database of 67,563 georeferenced occurrences collected during multiple fieldwork campaigns and complemented with records from museum collections available for the most part on https://heliconius-maps.github.io/ (accessed on November 2020; 114). We updated the taxonomic identity of records in agreement with the literature up to June 2021 (45, 46, but see Núñez et al. (47) for recent proposed taxonomic splits). This database covering 73 out of 77 species of the tribe (94.8 %) and 439 out of 457 subspecies (96.1 %) is available in <u>10.5281/zenodo.10906853</u>.

We employed the phylogeny of the tribe Heliconiini in Kozak et al. (45) encompassing 67 of the 77 recognized species (87 %) to estimate indices of phylogenetic diversity and evaluate niche convergence. However, we repeated the Bayesian estimations of divergence times between Heliconiini updating the secondary calibration points in accordance with recent estimates for Papilionoidea (31): Heliconiini-Acraeini (31.9-43.9 Mya); *Podotricha-Philaethria* (14.2-21.1 Mya), *Heliconius-Eueides* (11.6-20.3 Mya). We ran four independent analyses of 100 million cycles each in BEAST v2.6 resulting in divergence estimates in line with those generated previously based on the same alignment (45), as well as an independent estimate from genome-wide data (115). For Ithomiini, we used the phylogeny of Chazot et al. (48) that encompasses 339 of the 396 species (85.6 %). The divergence time used to bind the two tribes’ phylogenies was estimated at 84.49 My following Chazot et al. (31).

### 4.3 Species Distribution Modeling (SDM)

To predict spatial distributions, we performed Species Distribution Modeling (SDM) for each subspecies of Heliconiini independently. Modeling was carried out at subspecies level because many species are polymorphic, thus may belong to several phenotypic groups. The output of the SDM process was a single consensus model (ensemble model) for each subspecies that provides a proxy of probability of presence of each subspecies in each 30 km × 30 km grid cell (i.e., community).

As predictors of subspecies distributions, we used environmental variables that are relevant to butterfly ecology, according to the literature. Temperature and precipitation are known to influence the development of host plants for butterflies (116), while elevation (83, 117) and forest cover (118) are important factors shaping heliconiine butterfly distribution. We extracted annual mean temperature, mean diurnal range, annual precipitation levels, and precipitation seasonality from WorldClim bioclimatic variables dataset (v2.1 accessed 02/2021; 119), elevation from the SRTM Dataset (http://srtm.csi.cgiar.org/; v.4.1 accessed on 03/2019; 120), and forest cover from the Landsat Tree Cover Continuous Fields dataset (accessed on 03/2019; 121) aggregated at a quarter-degree cell resolution (i.e., pixel of ca. 30 km × 30 km).

We modelled each subspecies using three different algorithms (Random Forest, Gradient Tree Boosting and Artificial Neural Network) crossed with three independent sets of pseudo-absences and three spatially structured cross-validation blocks. We calculated the median of all models that passed our quality evaluation process to create an ensemble model for each subspecies. We cropped each subspecies’ predicted distribution to a relevant area according to its occurrences using a taxon-specific buffered alpha-hull mask. Finally, we merged ensemble models to acquire predicted distribution maps for species, mimetic groups, and Operational Mimicry Units (OMUs). OMUs are defined as all individuals of a unique species, including those spread among several subspecies, that belong to the same mimetic group (44). Despite the downstream analyses being carried out at the OMU level, for the sake of simplicity we used ‘co-mimetic species’ and ‘phenotypically similar species’ in the text to refer to the OMUs sharing the same phenotypic pattern.

All distribution maps are available in <u>10.5281/zenodo.10903661</u>. More details about the modelling process are available in the ODMAP form (122) in **Appendix 8**. Similar models were already performed for Ithomiini at the OMU-level in Doré et al. (44). These predictions were used to compare diversity patterns and investigate spatial associations between the two tribes.

### 4.4 Diversity indices

We computed a series of indices for each community (represented by 30 km × 30 km grid cell) and mapped them throughout the whole distribution range of the two tribes:

- Species and phenotypic richness, computed as the number of predicted species/phenotypic groups per community.
- Faith’s Phylogenetic Diversity (55), computed as the sum of branch lengths of the phylogenetic tree including all taxa found in a community.
- Mean species & phenotypic geographic rarity (123), computed as the weighted proportion of species or phenotypic group with small geographical ranges per community. These indices inform on the areas where species/phenotypic groups with the smallest ranges are concentrated.
- Mean phenotypic group size, computed as the mean number of species per phenotypic group within each community. This index provides insights on the degree of pattern convergence in the community as a high mean phenotypic group size implies a high number of species harboring the same wing patterns locally.
To compare the index values between Heliconiini and Ithomiini, we mapped them together with a two-dimensional color scale, scaled by the maximum of each index for each tribe, thus describing relative patterns of biodiversity (**Fig. 2**). Additionally, we computed spatial correlation tests across all communities for each index, between the two tribes (**Table S1** in **Appendix 4**), and between Heliconiini biodiversity patterns (**Table S2** in **Appendix 4**). We ran Spearman’s rank correlation tests using Clifford’s sample size correction to account for positive spatial autocorrelation across grid cell values (124). Lastly, we evaluated differences in spatial heterogeneity of biodiversity patterns between the two tribes using an asymptotic test to compare Coefficients of Variation (CV; 125).

### 4.5 Test for spatial association among co-mimetic species

To detect effects of phenotypic group membership, and thus of mutualistic interactions, on the spatial distribution of Heliconiini and Ithomiini, we investigated the degree of co-occurrence of phenotypically similar species (i.e., species in the same phenotypic group) across grid cells. We computed pairwise Bray-Curtis dissimilarities (126), an index that quantifies differences between the distribution of two entities, in our case between pairs of OMUs. Thus, we calculated the mean Bray-Curtis dissimilarity of all OMUs within phenotypic groups, globally and individually, representing the average degree of spatial co-occurrence of phenotypically similar units. To test the significance of these statistics, we used permutation tests under the null hypothesis that phenotypic group membership (i.e., wing color patterns) has no effect on co-occurrence. Therefore, for each permutation, we randomized the wing pattern between all OMUs to investigate whether phenotypically similar species co-occur more than expected at random, globally, and within each phenotypic group. As such, an observed BC lower than 95% of the null distribution of obtained values indicates a significant signal for spatial congruence. These analyses were performed for Ithomiini and Heliconiini independently and for pairs of phenotypically similar OMUs formed between the two tribes labeled as ‘inter-tribe’ in subsequent analyses (**Fig. S6** in **Appendix 5**; **Table S3** in **Appendix 6**).

### 4.6 Test for niche evolution among co-mimetic species

In order to investigate whether mimicry led to an evolutionary association between climatic niches and wing color patterns, we performed comparative phylogenetic analyses as was previously done for Ithomiini in Doré et al. (50). Climatic niche was described as the centroid of OMU occurrences using bioclimatic variables employed during niche modeling (i.e., annual mean temperature, mean diurnal range, annual precipitation levels, precipitation seasonality).

First, we fit multivariate neutral evolution models to explain the distribution of niche centroid values on the phylogeny. We compared AICc of a Brownian motion model with models implementing additional Pagel’s lambda or/and Pagel’s kappa parameters accounting respectively for presence of phylogenetic signal and punctuated evolution associated with cladogenesis (127, 128), to select for the best fitted option. At the end, we selected an evolution model with a Pagel’s lambda of 0.798.

We used mean climatic distances (MCD) computed as the pairwise Euclidean distances between niche centroids in the climatic space to estimate the similarity of climatic niches between pairs of OMUs. To test an effect of mimicry on climatic niche evolution, we simulated the evolution of the climatic niche under the selected neutral evolutionary model (n = 999) to obtain a null distribution for the mean MCD between phenotypically similar OMUs. As such, an observed MCD lower than 95% of null statistics indicates a significant signal for niche convergence (**Fig. S7** in **Appendix 5**; **Table S4** in **Appendix 6**).

## Data availability

All R scripts used to conduct the analyses and generate the figures are available on GitHub (https://github.com/EddiePerochon/Heliconiini_Diversity) with all generated files stored on <u>10.5281/zenodo.14765685</u>. Phenotypic classification, occurrences data, and maps of the distribution of subspecies, OMUs, species, and phenotypic groups produced and used in this study are available from Zenodo (Occurrences data: 10.5281/zenodo.10906853; Distribution maps: 10.5281/zenodo.10903661; Phenotypic classification: 10.5281/zenodo.10903197). All results reported in this article can be reproduced with the scripts and data provided.

## Author contributions

EP, ME and MD conceived the study and designed the analyses. NR, KK, WOM, BH, JM, JR & KW provided specimens and occurrence data. KK provided the phylogenetic trees. MD aggregated the database. EP curated the database. EP & MD wrote the R scripts. EP & MD carried out the analyses, produced maps, results, and figures. EP & MD led the manuscript writing. All authors contributed to the final manuscript.

## Acknowledgments

We gratefully thank all the researchers, field assistants, technicians and students who were involved in the collection of the occurrence data over the past decades. No specimens were collected within the scope of this research project. All georeferenced data were aggregated from previous research efforts; therefore, the authors declare no funding regarding data collection. EP’s research was funded by the Agence Nationale de la Recherche (ANR) under the grant CLEARWING (ANR-16-CE02-0012), and a Human Frontier Science Program grant (RGP0014/2016), both obtained by ME. MD acknowledges funding by the French Ministry of Research (MESRI). The authors certify that they have no affiliations with or involvement in any organization or entity with any financial interest, or non-financial interest in the subject matter or materials discussed in this article.

