## Supplementary Information - Appendices for "Müllerian mimicry in Neotropical butterflies: One mimicry ring to bring them all, and in the jungle bind them"

**This PDF file includes:**

- Supporting Information as Appendix 1 to 8.
- Figures S1 to S12.
- Tables S1 to S6.

### Appendix 1: Mimetic groups in Heliconiini and Ithomiini

We classified subspecies of Heliconiini butterflies into 38 phenotypic groups based on dorsal wing pattern similarity (**Fig. S1**) and representing ‘putative’ local mimicry rings, as in Doré et al. (1). Because those groups are based only on phenotypic similarity, members of such groups may not currently be involved in mutualistic interactions as they may not actually co-occur. If a significant signal of spatial co-occurrence within a phenotypic group is detected, it then qualifies as an ‘effective mimicry ring’, tentatively depicting true ecological interactions (2, 3).

We labeled each phenotypic group following the rationale of a taxonomic classification. For each group, we defined a ‘type’ taxon whose name was used to identify the phenotypic group and associated wing color pattern. The hierarchical criteria used to select the ‘type taxa’ were as follows:

- 1/ Avoiding the selection of a taxon whose name is associated with a polymorphic clade. This criterion aims to prevent confusion by forbidding the labeling of phenotypic groups with a name associated with multiple patterns. For instance, among taxa with large white bands on both hindwings and forewings, the most widely spread taxa are *Heliconius cydno cydno* and *Heliconius sapho sapho*. However, both *Heliconius cydno* and *Heliconius sapho* are polymorphic species. Therefore, we selected instead *Heliconius cydno chioneus* as the ‘type taxon’, and labeled the phenotypic group CHIONEUS, preventing any confusion potentially arising from a CYDNO or SAPHO group whose names refer to species with a diversity of wing patterns.
- 2/ Selecting the taxa with the broadest geographic range, and/or the most abundant, and/or the most widely known. The idea here is to favor the selection of the most widely recognized taxa among similar-looking taxa as the type. For instance, among taxa displaying a black background with a red band on the forewings, the most widespread and best-known taxon is *Heliconius erato hydara*, which is common throughout all the north of South America. Thus, we referred to this phenotypic group as HYDARA.
- 3/ Selecting the taxa with the oldest name, when multiple taxa are equivalent according to the previous criteria. For instance, *Heliconius pachinus* and *Heliconius hewitsoni* display similar triple white banded patterns and are found in similar areas across Central America. Since

*Heliconius pachinus* was named in 1871, while *Heliconius hewitsoni* was named in 1875, we selected *Heliconius pachinus* as the 'type taxon' and labeled the phenotypic group PACHINUS.

Moreover, to ensure robustness of our results to alternative choices of phenotypic classifications, we designed higher-level phenotypic groups that encompass multiple lower-level groups using broader color pattern features to define phenotypic similarity. We ran all analyses using the two most extreme choices for the classification: the most 'split' with 38 phenotypic groups as showed in the main text, and the most 'lumped' with 20 phenotypic groups. This design ensures that any intermediate choice in the phenotypic classification would lead to similar results, as long as results of the two extreme options lead to similar conclusions.

The comprehensive phenotypic classification of heliconiine subspecies is available in the online archive, alongside pictures of the 'type taxon' of each phenotypic group (see 10.5281/zenodo.10903197). This classification includes a description of key features used to identify and discriminate between the phenotypic groups for both, the most 'split' classification used in the main text, and the most 'lumped' classification used for sensitivity analyses (see **Appendix 7**).

For Ithomiini, we used the currently accepted classification of mimicry patterns (**Fig. S2**, 4), built using a similar rationale of phenotypic similarity, independent from geographical distributions. Then, we matched the identity of phenotypic groups associated with a pattern represented in the two tribes and labeled them as inter-tribe phenotypic groups (dash frames in **Figs. S1 & S2**). Because ithomiine phenotypic groups were defined prior to heliconiine phenotypic groups, the inter-tribe phenotypic groups retained their initial name based on an ithomiine 'type taxon' (see 10.5281/zenodo.5497876).

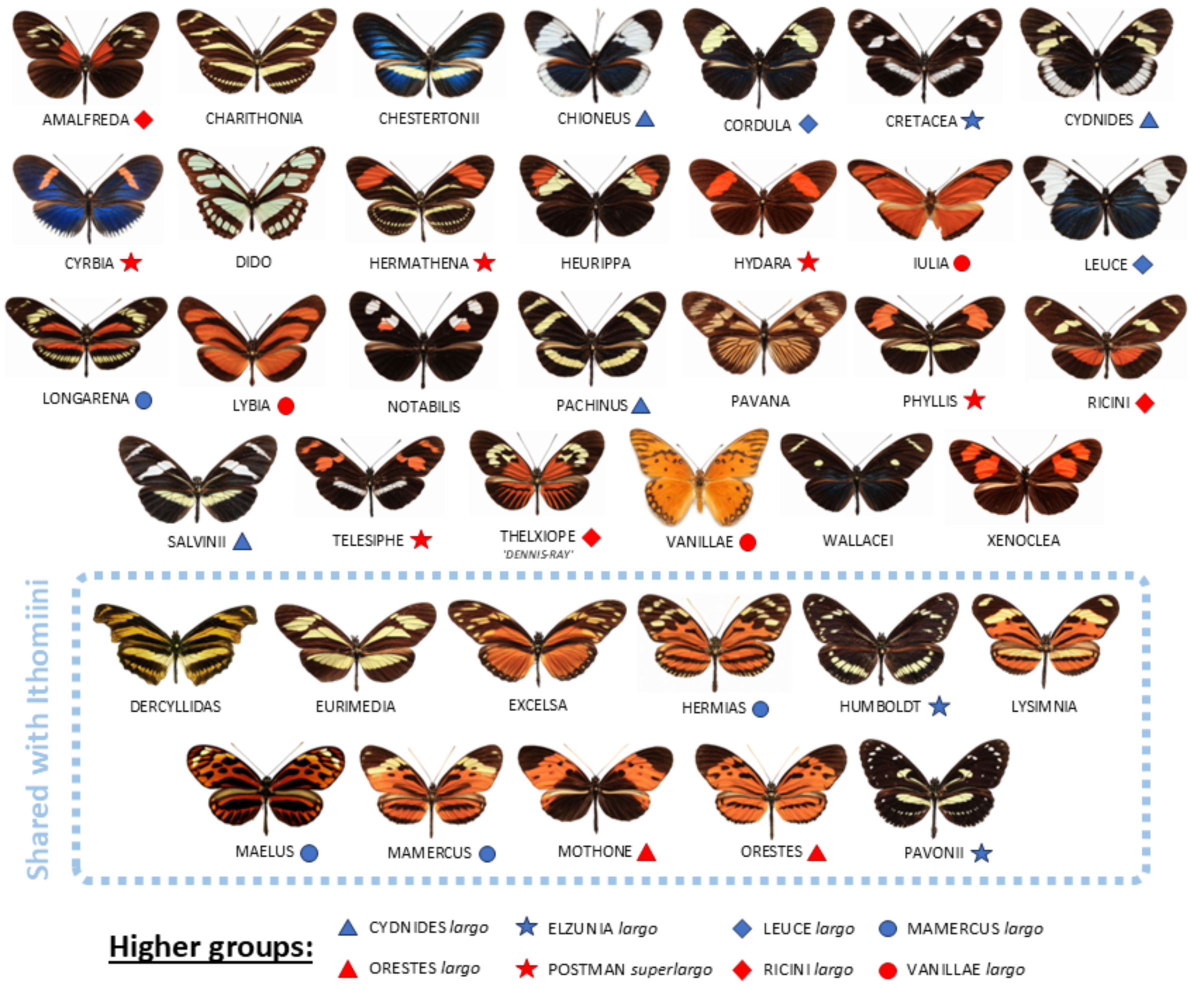


**Figure S1: Phenotypic groups in Heliconiini.** ‘Type taxon’ for each heliconiine phenotypic group representing the color pattern shared by all members of such groups used in this study. Groups are designed based on phenotypic similarity in shape, patterns, and colors. Labels of the groups were chosen to reflect the name of the selected ‘type taxon’. Groups shown within the dash frame are inter-tribe phenotypic groups shared with ithomiine butterflies. They are labeled after ithomiine type taxon (1, 4). Groups shown here correspond to the most ‘split’ classification used in analyses presented in the main text. Higher groups encompassing multiple lower-level groups are aggregated according to the symbols. They were used to carry out sensitivity analyses using the most ‘lumped’ classification as an extreme alternative (See **Appendix 7**). Photo credits: C. Jiggins.

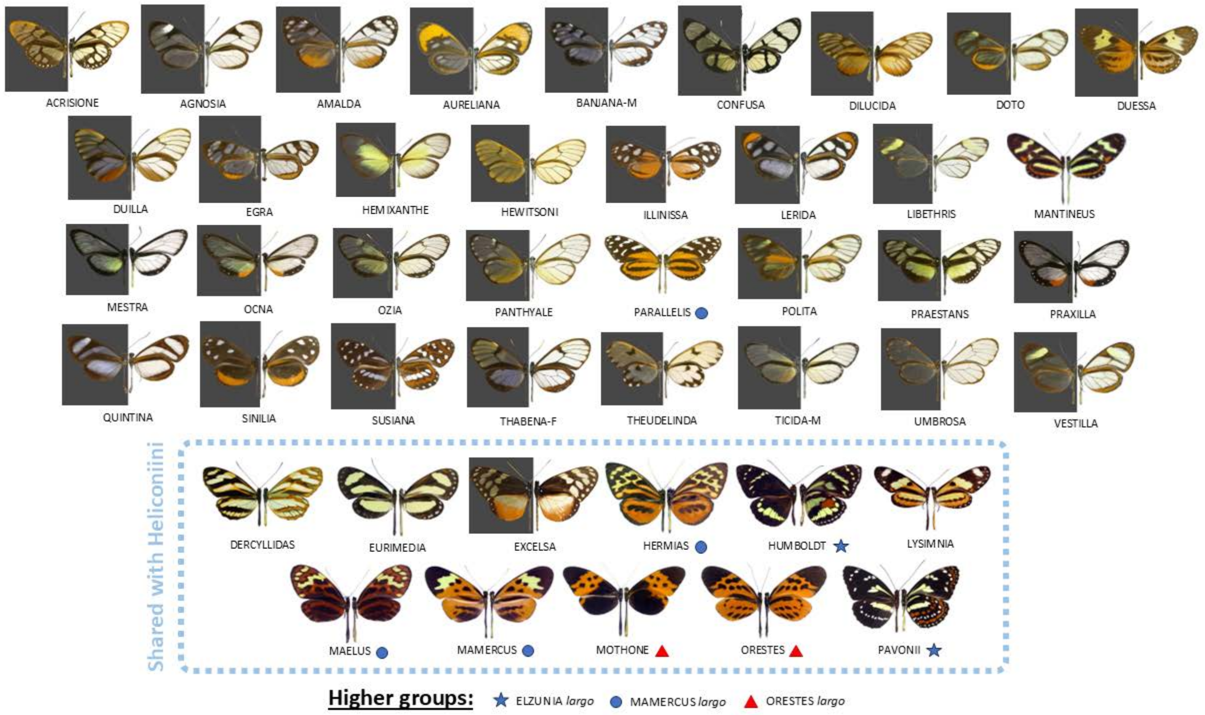


**Figure S2: Phenotypic groups in Ithomiini.** Type taxon of each ithomiine phenotypic group representing the color pattern shared by all members of such groups. Groups were designed in a previous study based on phenotypic similarity in shape, patterns, and colors (4). Groups shown within the dash frame are inter-tribe phenotypic groups shared with heliconiine butterflies. Shared groups shown here correspond to the most 'split' classification used in analyses presented in the main text. Higher groups encompassing multiple lower-level groups are aggregated according to the symbols. They were used to carry out sensitivity analyses for inter-tribe groups using the most 'lumped' classification as an extreme alternative (See **Appendix 7**). Figure adapted from Doré et al. (1). Photo credits: K. Willmott.

#### Appendix 2: Map of bioregions

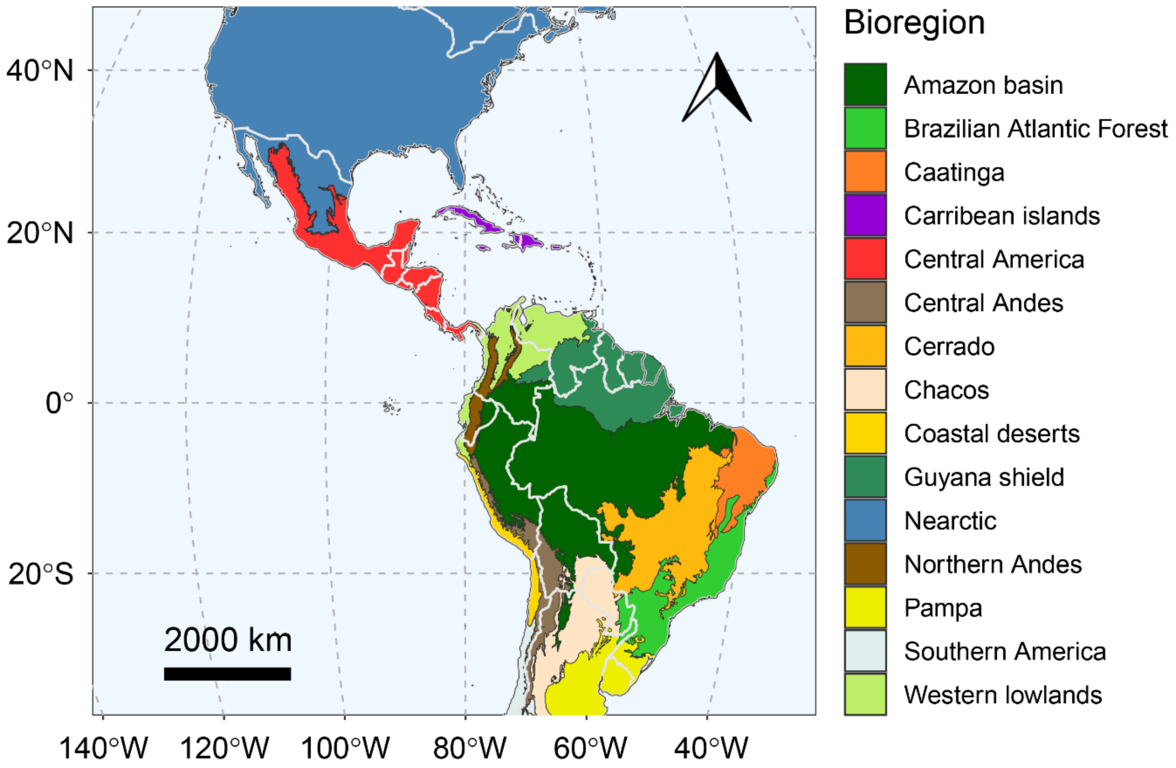


**Figure S4: Map of bioregions.** Bioregions used throughout the study to describe biodiversity patterns are based on the aggregation of provinces defined in Morrone et al. (5). White lines represent geopolitical boundaries between countries.

### Appendix 3: Heatmaps of biodiversity patterns

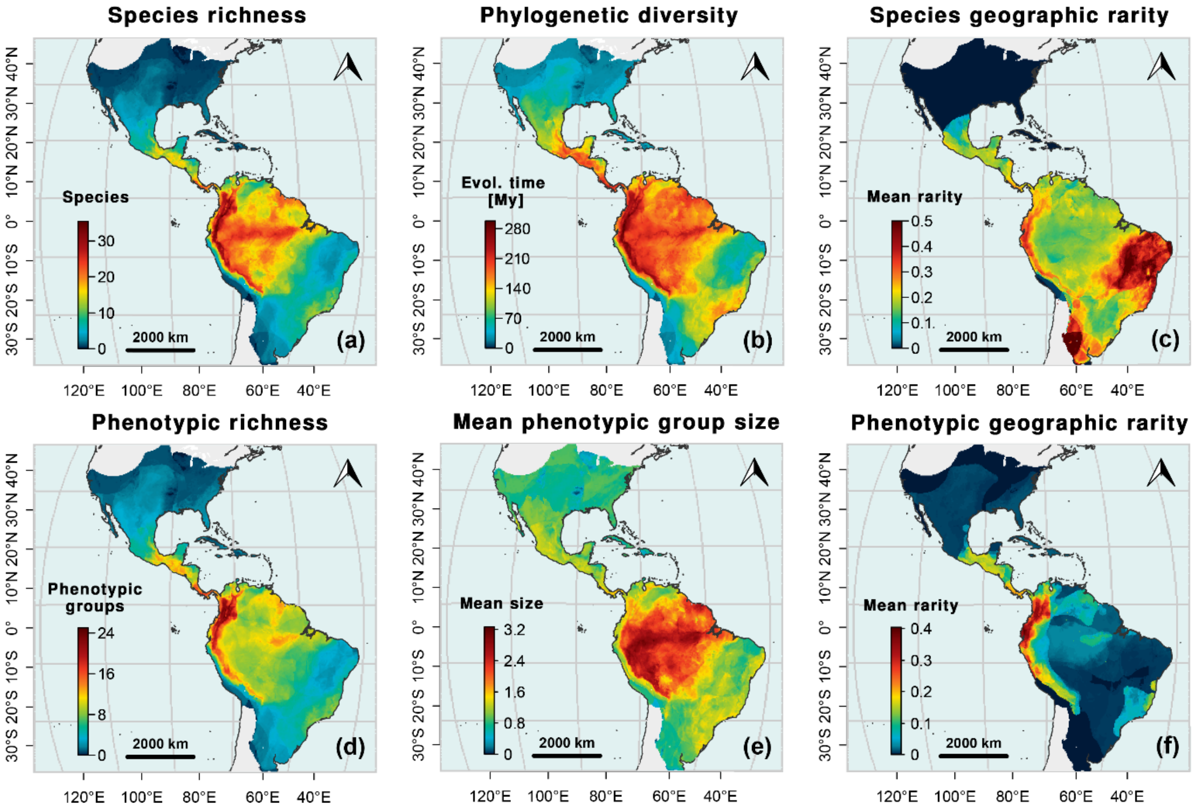


**Figure S4: Heliconiini biodiversity patterns. (a)** Species richness. **(b)** Faith's Phylogenetic Diversity Faith's Phylogenetic Diversity (6). **(c)** Mean species geographic rarity. Rarity index based on species ranges. **(d)** Phenotypic richness (i.e., number of local phenotypic groups). **(e)** Mean phenotypic geographic rarity. Rarity index based on phenotypic group ranges. **(f)** Mean phenotypic group size (i.e., mean number of species per local phenotypic groups).

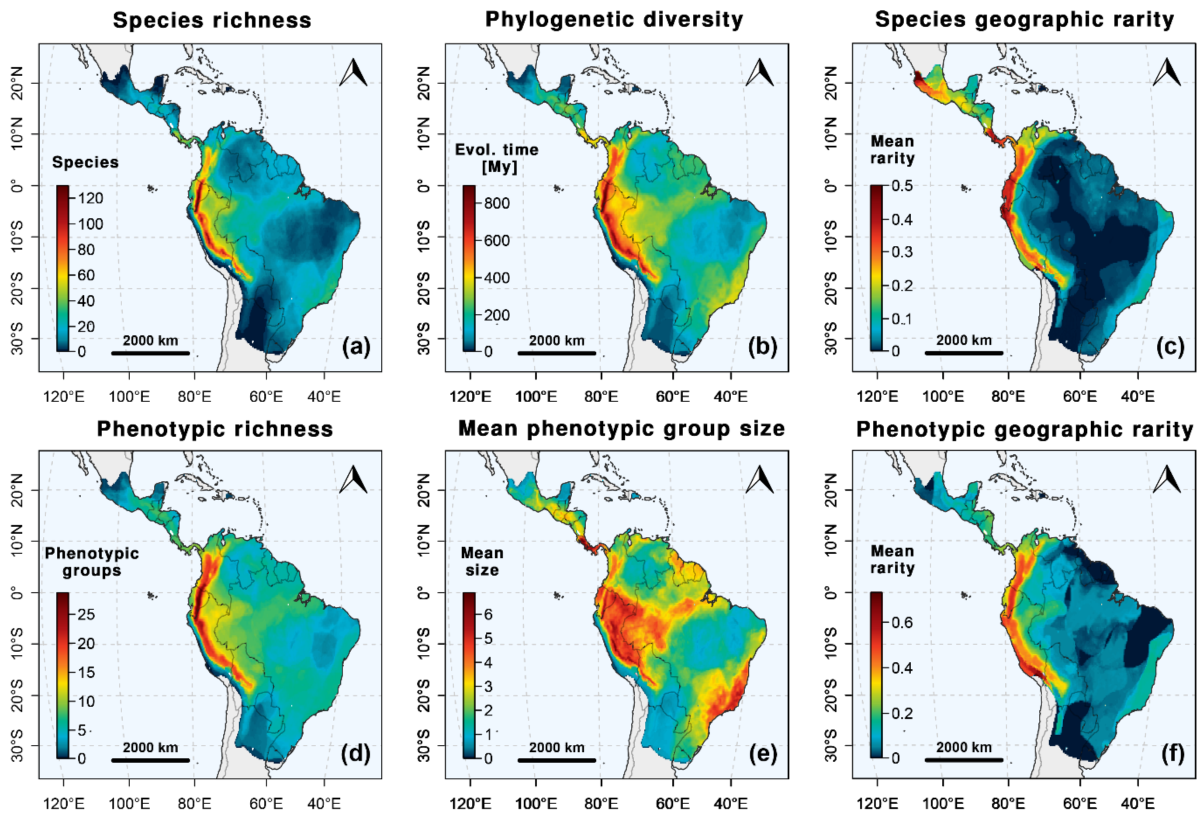


**Figure S5: Ithomiini biodiversity patterns.** Modified from Figure 3 in Doré et al. (4). **(a)** Species richness. **(b)** Faith's Phylogenetic Diversity (6). **(c)** Mean species geographic rarity. Rarity index based on species ranges. **(d)** Phenotypic richness (i.e., number of local phenotypic groups). **(e)** Mean phenotypic geographic rarity. Rarity index based on phenotypic group ranges. **(f)** Mean phenotypic group size (i.e., mean number of species per local phenotypic groups).

### Appendix 4: Tables for spatial correlation tests

**Table S1: Tests for spatial correlation of biodiversity indices between Heliconiini and Ithomiini tribes.** N = 18,006 grid cells. Rho = Spearman's rank correlation coefficient. Df adjusted with Clifford's correction (7). Q95 = Threshold value for $\alpha = 0.05$. Significance levels: * = $p < 0.05$; ** = $p < 0.01$; *** = $p < 0.001$.

| Index | Rho | T-stat | Adjusted df | Q95 | p-value |
|---|---|---|---|---|---|
| Species richness | 0.771 | 7.91 | 42.7 | 1.681 | < 0.001 *** |
| Phylogenetic diversity | 0.779 | 8.34 | 45.1 | 1.679 | < 0.001 *** |
| Mean species geographic rarity | -0.042 | -0.34 | 64.7 | 1.669 | 0.632 |
| Phenotypic richness | 0.762 | 7.60 | 41.8 | 1.682 | < 0.001 *** |
| Mean phenotypic group size | 0.626 | 5.87 | 53.5 | 1.674 | < 0.001 *** |
| Mean phenotypic geographic rarity | 0.625 | 5.71 | 50.9 | 1.675 | < 0.001 *** |

**Table S2: Tests for spatial correlation between Heliconiini species richness and other biodiversity indices.** N = 26,539 grid cells. Rho = Spearman's rank correlation coefficient. Df adjusted with Clifford's correction (7). Q95 = Threshold value for $\alpha = 0.05$. Significance levels: * = $p < 0.05$; ** = $p < 0.01$; *** = $p < 0.001$.

| Index | Rho | T-stat | Adjusted df | Q95 | p-value |
|---|---|---|---|---|---|
| Phylogenetic diversity | 0.987 | 24.2 | 15.5 | 1.749 | < 0.001 *** |
| Mean species geographic rarity | 0.448 | 2.0 | 15.9 | 1.747 | 0.032 * |
| Phenotypic richness | 0.978 | 18.0 | 14.7 | 1.755 | < 0.001 *** |
| Mean phenotypic group size | 0.909 | 8.4 | 14.8 | 1.754 | < 0.001 *** |
| Mean phenotypic geographic rarity | 0.72 | 5.5 | 27.9 | 1.701 | < 0.001 *** |

### Appendix 5: Global tests for spatial congruence and niche convergence

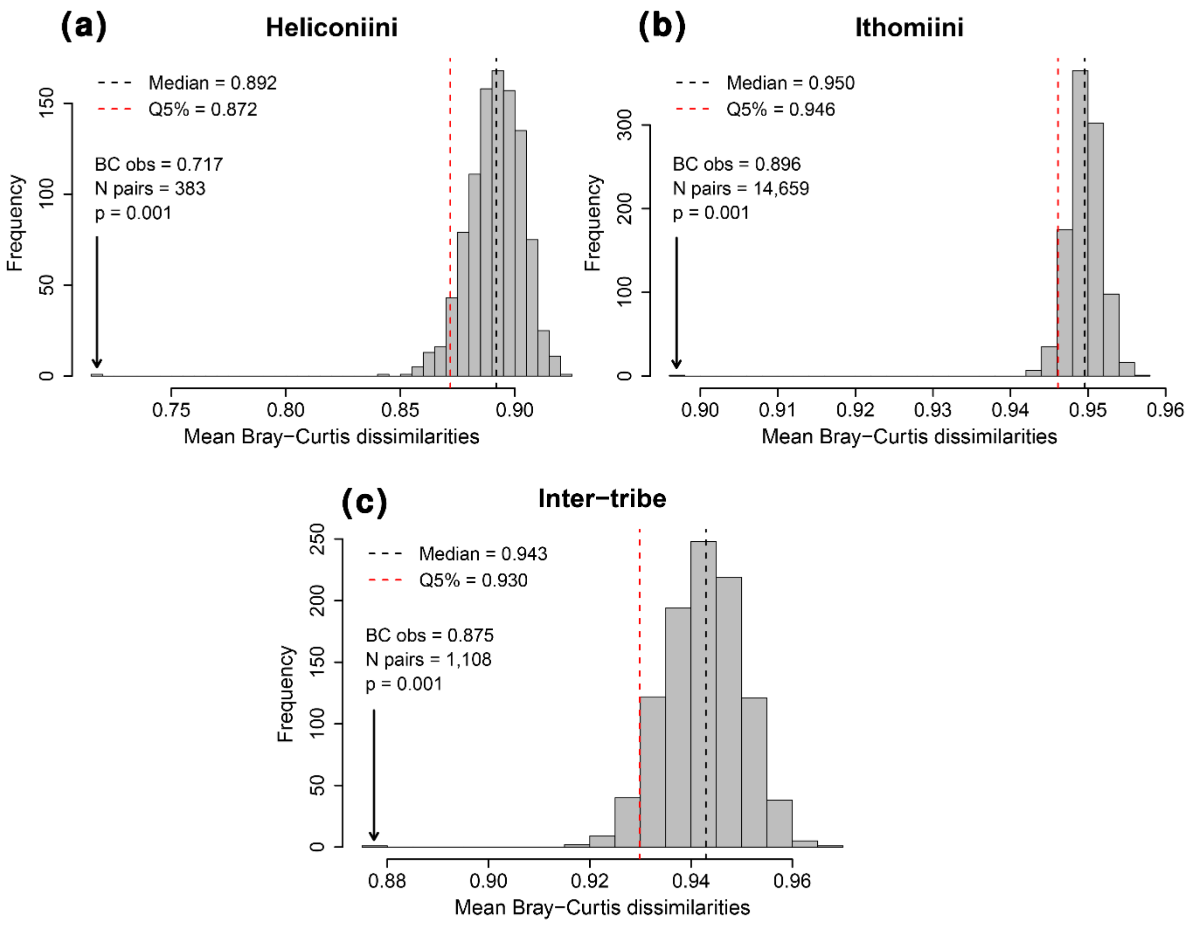


**Figure S6: Null distributions for spatial congruence tests within phenotypic groups. (a)** Heliconiini. **(b)** Ithomiini adapted from Doré et al. (1). **(c)** Inter-tribe phenotypic groups. Permutation tests with 1000 randomizations. Bray-Curtis (BC) indices quantify the degree of dissimilarity in spatial distribution between pairs of OMUs. An observed mean BC value lower than the 5% quantile (Q5%) of the null distribution supports the overall spatial congruence of phenotypically similar species.

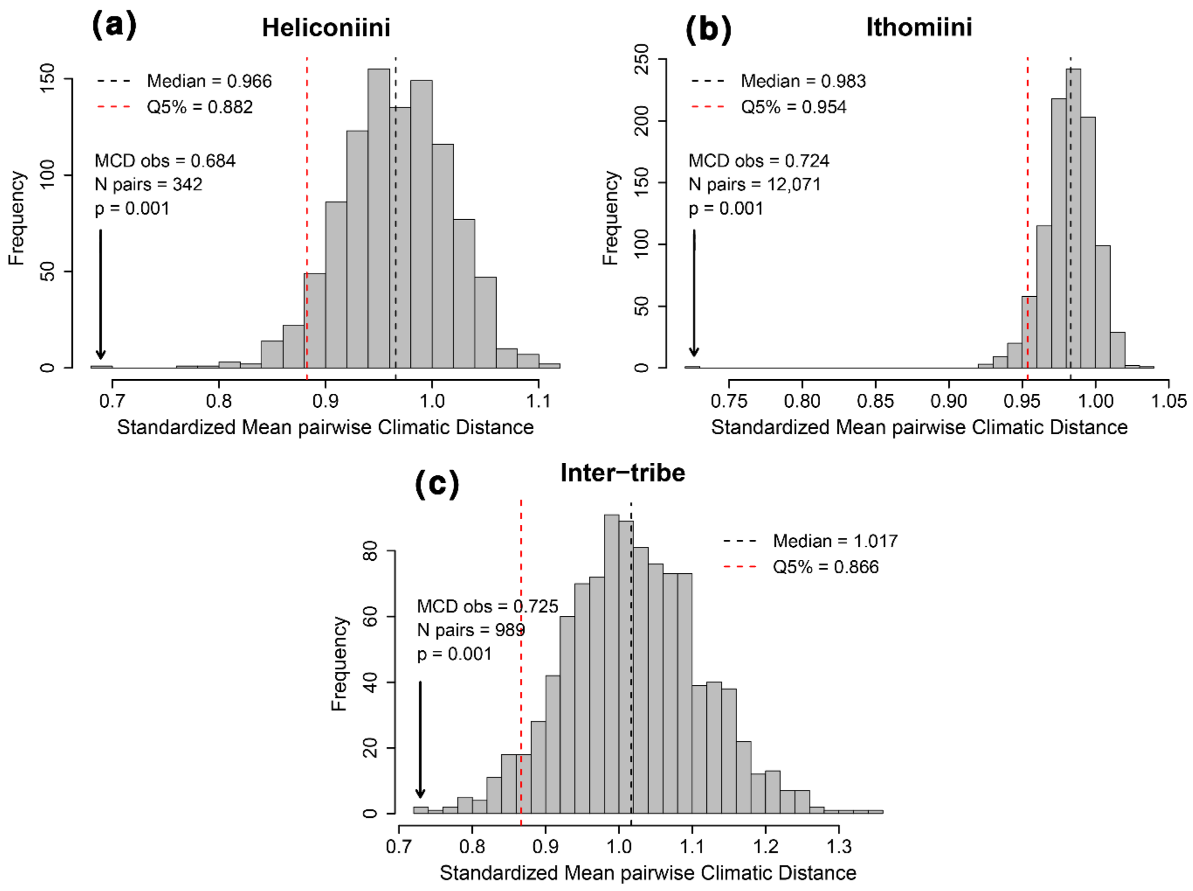


**Figure S7: Null distributions for niche convergence tests within phenotypic groups. (a)** Heliconiini **(b)** Ithomiini adapted from Doré et al. (1) **(c)** Inter-tribe phenotypic groups. Phylogenetic comparative tests with 1000 simulations of neutral niche evolution (Brownian Motion model with Pagel's lambda = 0.798). Standardized Mean pairwise Climatic Distances (MCD) quantify the degree of dissimilarity in niche centroids between pairs of OMUs. An observed mean MCD value lower than the 5% quantile (Q5%) of the null distribution supports the overall niche convergence of phenotypically similar species.

### Appendix 6: Phenotypic group-level tests for spatial congruence and niche convergence

**Table S3: Tests for spatial congruence within each phenotypic group.** Bray-Curtis (BC) indices quantify the degree of dissimilarity in spatial distribution between pairs of OMUs within a phenotypic group. An observed mean BC value lower than the 5% quantile (Q5%) of the null distribution supports the spatial congruence of phenotypically similar species within a phenotypic group. Tests for inter-tribe phenotypic groups are carried out between pairs of heliconiine vs. ithomiine OMUs. $BC_{obs}$ = Observed mean pairwise BC. $BC_{Q50}$ = Median of the null distribution. $BC_{Q5}$ = Quantile 5% of the null distribution = statistical significance threshold for $\alpha = 0.05$. P-value for the spatial congruence test is based on the rank of the $BC_{obs}$ among the null distribution obtained through random permutation of phenotypic group membership (N = 1000): * = $p < 0.05$; ** = $p < 0.01$; *** = $p \leq 0.001$.

| PHENOTYPIC GROUP | N units | N pairs | $BC_{obs}$ | $BC_{Q50}$ | $BC_{Q5}$ | p-value |
|---|---|---|---|---|---|---|
| **Heliconiini groups** | | | | | | |
| AMALFREDA | 10 | 45 | 0.636 | 0.896 | 0.815 | 0.001*** |
| CHARITHONIA | 1 | 0 | | | | |
| CHESTERTONII | 1 | 0 | | | | |
| CHIONEUS | 2 | 1 | 0.437 | 0.977 | 0.496 | 0.04* |
| CORDULA | 3 | 3 | 0.999 | 0.924 | 0.691 | 0.883 |
| CRETACEA | 1 | 0 | | | | |
| CYDNIDES | 2 | 1 | 0.605 | 0.981 | 0.493 | 0.09 |
| CYRBIA | 2 | 1 | 0.172 | 0.968 | 0.463 | 0.002** |
| DERCYLLIDAS | 1 | 0 | | | | |
| DIDO | 6 | 15 | 0.897 | 0.903 | 0.785 | 0.462 |
| EURIMEDIA | 1 | 0 | | | | |
| EXCELSA | 4 | 6 | 0.679 | 0.906 | 0.737 | 0.025* |
| HERMATHENA | 1 | 0 | | | | |
| HERMIAS | 7 | 21 | 0.835 | 0.9 | 0.797 | 0.143 |
| HEURIPPA | 1 | 0 | | | | |
| HUMBOLDT | 2 | 1 | 0.411 | 0.977 | 0.566 | 0.019* |
| HYDARA | 3 | 3 | 0.716 | 0.921 | 0.686 | 0.078 |
| IULIA | 4 | 6 | 0.297 | 0.907 | 0.727 | 0.001*** |
| LEUCE | 4 | 6 | 0.789 | 0.904 | 0.697 | 0.133 |
| LONGARENA | 1 | 0 | | | | |
| LYBIA | 5 | 10 | 0.93 | 0.909 | 0.752 | 0.648 |
| LYSIMNIA | 3 | 3 | 0.71 | 0.916 | 0.697 | 0.06 |
| MAELUS | 3 | 3 | 0.661 | 0.915 | 0.665 | 0.048* |
| MAMERCUS | 7 | 21 | 0.811 | 0.897 | 0.788 | 0.103 |
| MOTHONE | 1 | 0 | | | | |
| NOTABILIS | 2 | 1 | 0.182 | 0.981 | 0.511 | 0.002** |
| ORESTES | 6 | 15 | 0.611 | 0.906 | 0.768 | 0.002** |
| PACHINUS | 3 | 3 | 0.251 | 0.918 | 0.702 | 0.001*** |
| PAVANA | 3 | 3 | 0.971 | 0.927 | 0.707 | 0.68 |
| PAVONII | 3 | 3 | 0.746 | 0.919 | 0.685 | 0.096 |

| | | | | | | |
|---|---|---|---|---|---|---|
| PHYLLIS | 4 | 6 | 0.81 | 0.913 | 0.736 | 0.141 |
| RICINI | 7 | 21 | 0.93 | 0.898 | 0.779 | 0.759 |
| SALVINII | 2 | 1 | 1 | 0.967 | 0.512 | 1 |
| TELESIPHE | 2 | 1 | 0.148 | 0.981 | 0.521 | 0.002** |
| THELXIOPE | 17 | 136 | 0.686 | 0.895 | 0.834 | 0.001*** |
| VANILLAE | 5 | 10 | 0.843 | 0.9 | 0.751 | 0.214 |
| WALLACEI | 9 | 36 | 0.67 | 0.9 | 0.798 | 0.003** |
| XENOCLEA | 2 | 1 | 0.395 | 0.966 | 0.441 | 0.037* |
| **Inter-tribe groups** | | | | | | |
| DERCYLLIDAS | 2 | 1 | 0.261 | 1 | 0.675 | 0.002** |
| EURIMEDIA | 36 | 35 | 0.979 | 0.942 | 0.872 | 0.798 |
| EXCELSA | 22 | 72 | 0.805 | 0.943 | 0.894 | 0.001*** |
| HERMIAS | 60 | 371 | 0.9 | 0.943 | 0.917 | 0.006** |
| HUMBOLDT | 3 | 2 | 0.508 | 0.991 | 0.738 | 0.006** |
| LYSIMNIA | 8 | 15 | 0.728 | 0.95 | 0.867 | 0.001*** |
| MAELUS | 19 | 48 | 0.643 | 0.945 | 0.894 | 0.001*** |
| MAMERCUS | 71 | 448 | 0.927 | 0.944 | 0.918 | 0.138 |
| MOTHONE | 15 | 14 | 0.731 | 0.95 | 0.863 | 0.001*** |
| ORESTES | 22 | 96 | 0.742 | 0.944 | 0.901 | 0.001*** |
| PAVONII | 5 | 6 | 0.653 | 0.952 | 0.819 | 0.001*** |

**Table S4: Tests for niche convergence within each phenotypic group.** Standardized Mean pairwise Climatic Distances (MCD) quantify the degree of dissimilarity in niche centroids between pairs of OMUs. An observed mean MCD value lower than the 5% quantile (Q5%) of the null distribution obtained through simulation of the neutral evolution of the niche supports the niche convergence of phenotypically similar species within a phenotypic group. Tests for inter-tribe phenotypic groups are carried out between pairs of heliconiine vs. ithomiine OMUs. $MCD_{obs}$ = Observed MCD. $MCD_{Q50}$ = Median of the null distribution. $MCD_{Q5}$ = Quantile 5% of the null distribution = statistical significance threshold for α = 0.05. P-value for the spatial congruence test is based on the rank of the $MCD_{obs}$ among the null distribution obtained through neutral evolution of the niche (N = 1000): * = p < 0.05; ** = p < 0.01; *** = p ≤ 0.001.

| PHENOTYPIC GROUP | N units | N pairs | $MCD_{obs}$ | $MCD_{Q50}$ | $MCD_{Q5}$ | p-value |
|---|---|---|---|---|---|---|
| **Heliconiini groups** | | | | | | |
| AMALFREDA | 9 | 36 | 0.339 | 0.993 | 0.721 | 0.001*** |
| CHARITHONIA | 1 | 0 | | | | |
| CHESTERTONII | 1 | 0 | | | | |
| CHIONEUS | 2 | 1 | 0.252 | 0.901 | 0.268 | 0.044* |
| CORDULA | 3 | 3 | 1.641 | 0.88 | 0.399 | 0.971 |
| CRETACEA | 1 | 0 | | | | |
| CYDNIDES | 2 | 1 | 0.78 | 1.109 | 0.275 | 0.306 |
| CYRBIA | 2 | 1 | 0.284 | 1.102 | 0.303 | 0.041* |
| DIDO | 5 | 10 | 0.78 | 0.745 | 0.438 | 0.558 |
| EURIMEDIA | 1 | 0 | | | | |

| | | | | | | |
|---|---|---|---|---|---|---|
| EXCELSA | 4 | 6 | 1.908 | 1.019 | 0.544 | 0.994 |
| HERMATHENA | 1 | 0 | | | | |
| HERMIAS | 7 | 21 | 0.62 | 1.082 | 0.721 | 0.015* |
| HEURIPPA | 1 | 0 | | | | |
| HUMBOLDT | 2 | 1 | 1.177 | 0.957 | 0.277 | 0.661 |
| HYDARA | 3 | 3 | 0.385 | 1.062 | 0.491 | 0.021* |
| IULIA | 4 | 6 | 0.3 | 1.024 | 0.548 | 0.003** |
| JUDITH | 1 | 0 | | | | |
| LEUCE | 4 | 6 | 0.827 | 1.021 | 0.58 | 0.257 |
| LONGARENA | 1 | 0 | | | | |
| LYBIA | 3 | 3 | 0.896 | 0.844 | 0.393 | 0.575 |
| LYSIMNIA | 3 | 3 | 0.366 | 0.945 | 0.46 | 0.027* |
| MAELUS | 3 | 3 | 0.807 | 0.828 | 0.376 | 0.479 |
| MAMERCUS | 7 | 21 | 1.079 | 1.067 | 0.721 | 0.519 |
| MOTHONE | 1 | 0 | | | | |
| NOTABILIS | 2 | 1 | 0.503 | 1.137 | 0.341 | 0.115 |
| ORESTES | 6 | 15 | 0.522 | 1.068 | 0.699 | 0.002** |
| PACHINUS | 3 | 3 | 0.245 | 0.791 | 0.358 | 0.013* |
| PAVANA | 3 | 3 | 0.644 | 0.796 | 0.362 | 0.297 |
| PAVONII | 3 | 3 | 1.713 | 0.88 | 0.407 | 0.978 |
| PHYLLIS | 4 | 6 | 0.873 | 0.95 | 0.531 | 0.394 |
| RICINI | 7 | 21 | 1.245 | 0.882 | 0.606 | 0.974 |
| SALVINII | 2 | 1 | 1.438 | 0.907 | 0.239 | 0.844 |
| TELESIPHE | 2 | 1 | 0.112 | 0.977 | 0.269 | 0.011* |
| THELXIOPE | 16 | 120 | 0.532 | 0.962 | 0.778 | 0.001*** |
| VANILLAE | 4 | 6 | 0.904 | 0.863 | 0.472 | 0.56 |
| WALLACEI | 9 | 36 | 0.749 | 0.846 | 0.606 | 0.253 |
| XENOCLEA | 2 | 1 | 0.057 | 1.111 | 0.318 | 0.002** |
| **Inter-tribe groups** | | | | | | |
| EURIMEDIA | 34 | 33 | 0.679 | 0.935 | 0.612 | 0.109 |
| EXCELSA | 21 | 68 | 1.036 | 0.995 | 0.76 | 0.609 |
| HERMIAS | 54 | 329 | 0.647 | 1.018 | 0.827 | 0.001*** |
| HUMBOLDT | 3 | 2 | 0.561 | 0.952 | 0.412 | 0.136 |
| LYSIMNIA | 7 | 12 | 0.341 | 0.972 | 0.648 | 0.001*** |
| MAELUS | 18 | 45 | 0.525 | 0.974 | 0.703 | 0.001*** |
| MAMERCUS | 63 | 392 | 0.837 | 1.028 | 0.838 | 0.05* |
| MOTHONE | 13 | 12 | 0.448 | 0.972 | 0.612 | 0.003** |
| ORESTES | 21 | 90 | 0.471 | 1.028 | 0.788 | 0.001*** |
| PAVONII | 5 | 6 | 1.09 | 0.945 | 0.517 | 0.702 |

### Appendix 7: Sensitivity analyses with ‘lumped’ classification

To ensure robustness of our results to alternative choices of phenotypic classifications, we also ran all analyses on a classification that encompasses all the highest level (i.e., most ‘lumped’) phenotypic groups. Therefore, analyses have been carried out for the two most extreme choices for the classification: the most ‘split’ with 38 phenotypic groups as shown in the main text, and the most ‘lumped’ with 20 phenotypic groups as presented below. This design ensures that any intermediate choice in the phenotypic classification would lead to similar results, as long as results of the two extreme options lead to similar conclusions.

Our sensitivity analyses showed that that diversity indices of phenotypic richness, mean group size and mean geographic rarity (**Fig. S8.d-f**) displayed qualitatively similar patterns to what was obtained under the most ‘split’ classification used for the main analyses (**Fig. S1.d-f**). Since groups encompass more taxa in the ‘lumped classification’, differences reside in the absolute number of phenotypic groups and mean size of local groups that range up to 13 local groups in the Northern Andes and 5.6 taxa per groups in the Amazon Basin with the ‘lumped’ classification (**Fig. S1.d-e**), compared to 24 local groups and 3.2 taxa per groups with the ‘split’ classification.

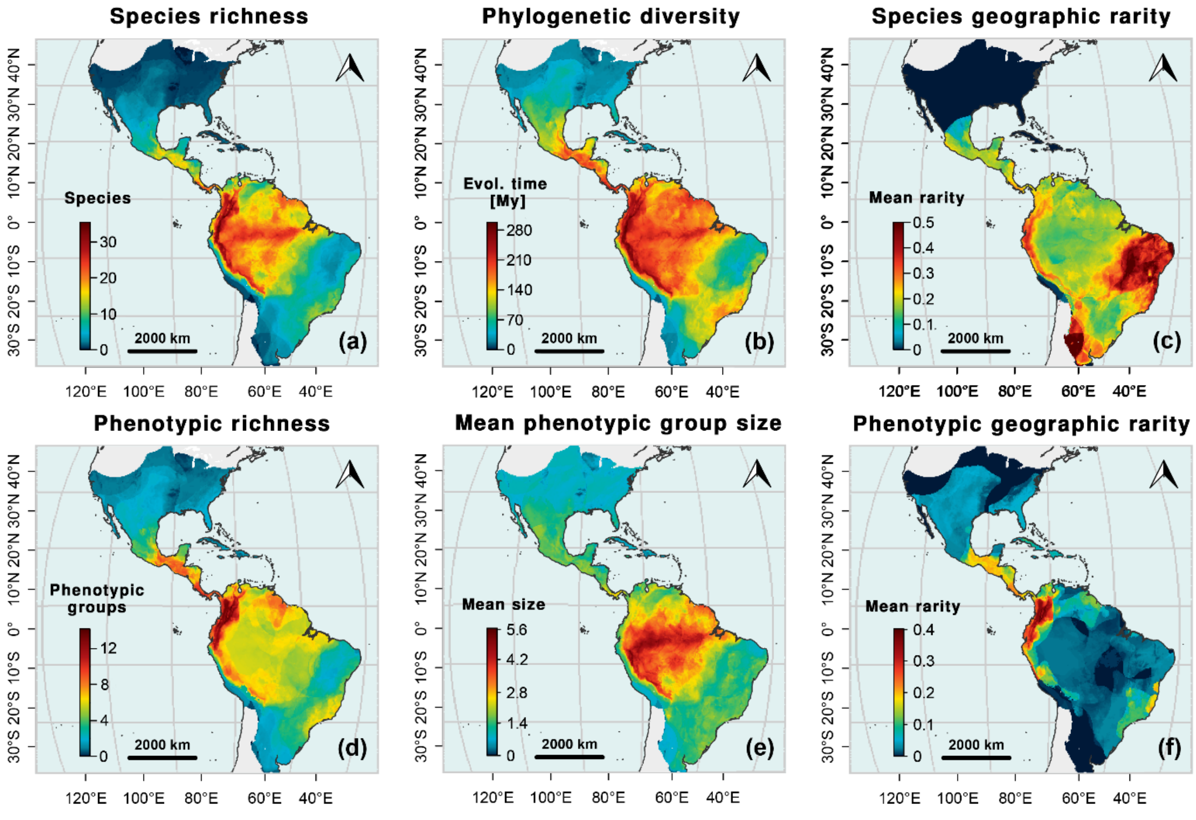

**Figure S8: Heliconiini biodiversity patterns using the 'lumped' classification. (a)** Species richness. **(b)** Faith's Phylogenetic Diversity (6). **(c)** Mean species geographic rarity. Rarity index based on species ranges. **(d)** Phenotypic richness (i.e., number of local phenotypic groups). **(e)** Mean phenotypic geographic rarity. Rarity index based on phenotypic group ranges. **(f)** Mean phenotypic group size (i.e., mean number of species per local phenotypic groups).

Patterns of co-occurrence in Heliconiine butterflies were supported indifferently with the 'split' classification (**Fig S6.a**; Permutation test: $BC_{obs}$ = 0.717, $BC_{null}$ Q5% = 0.872, p ≤ 0.001), or the 'lumped' classification (**Fig S9.a**; Permutation test: $BC_{obs}$ = 0.774, $BC_{null}$ Q5% = 0.847, p ≤ 0.001). Similarly, we found a signal for significance co-occurrence between pairs of Heliconiini and Ithomiini taxa (i.e., inter-tribes) sharing similar phenotypes for both the 'split' classification (**Fig S6.c**; Permutation test: $BC_{obs}$ = 0.875, $BC_{null}$ Q5% = 0.930, p ≤ 0.001) and the 'lumped' classification (**Fig S9.b**; Permutation test: $BC_{obs}$ = 0.839, $BC_{null}$ Q5% = 0.921, p ≤ 0.001). At group-level, we detected a significant signal for spatial congruence in 6 out of 15 (40.0%) 'lumped' phenotypic groups of heliconiines (**Table S5**). This proportion rises to 6 out of 7 (85.7%) 'lumped' groups for inter-tribe phenotypic groups shared between the heliconiine and ithomiine butterflies (**Table S5**).

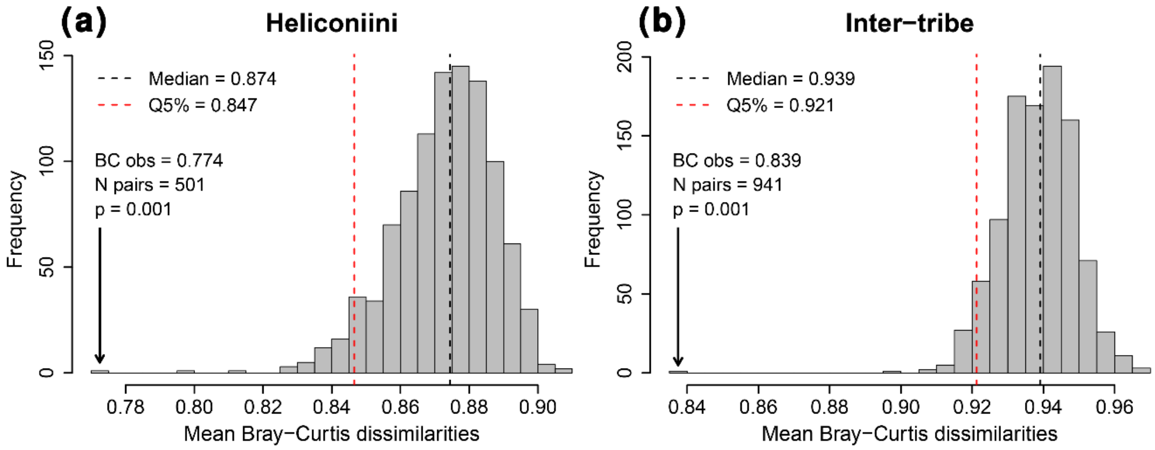


**Figure S9: Null distributions for spatial congruence tests within phenotypic groups using the 'lumped' classification. (a)** Heliconiini. **(b)** Inter-tribe phenotypic groups. Permutation tests with 1000 randomizations. Bray-Curtis (BC) indices quantify the degree of dissimilarity in spatial distribution between pairs of OMUs. An observed mean BC value lower than the 5% quantile (Q5%) of the null distribution supports the overall spatial congruence of phenotypically similar species.

**Table S5: Tests for spatial congruence within each phenotypic group using the 'lumped' classification.** Bray-Curtis (BC) indices quantify the degree of dissimilarity in spatial distribution between pairs of OMUs within a phenotypic group. An observed mean BC value lower than the 5% quantile (Q5%) of the null distribution supports the spatial congruence of phenotypically similar species within a phenotypic group. Tests for inter-tribe phenotypic groups are carried out between pairs of heliconiine vs. ithomiine OMUs. $BC_{obs}$ = Observed mean pairwise BC. $BC_{Q50}$ = Median of the null distribution. $BC_{Q5}$ = Quantile 5% of the null distribution = statistical significance threshold for $\alpha = 0.05$. P-value for the spatial congruence test is based on the rank of the $BC_{obs}$ among the null distribution obtained through random permutation of phenotypic group membership (N = 1000): * = p < 0.05; ** = p < 0.01; *** = p ≤ 0.001.

| PHENOTYPIC GROUP | N units | N pairs | $BC_{obs}$ | $BC_{Q50}$ | $BC_{Q5}$ | p-value |
|---|---|---|---|---|---|---|
| **Heliconiini groups** | | | | | | |
| CHARITHONIA | 1 | 0 | | | | |
| CHESTERTONII | 1 | 0 | | | | |
| CYDNIDES_largo | 7 | 21 | 0.755 | 0.883 | 0.747 | 0.063 |
| DERCYLLIDAS | 1 | 0 | | | | |
| DIDO | 6 | 15 | 0.897 | 0.89 | 0.744 | 0.552 |
| ELZUNIA_largo | 5 | 10 | 0.799 | 0.893 | 0.731 | 0.145 |
| EURIMEDIA | 1 | 0 | | | | |
| EXCELSA | 4 | 6 | 0.679 | 0.892 | 0.686 | 0.048* |
| HEURIPPA | 1 | 0 | | | | |
| LEUCE_largo | 5 | 10 | 0.77 | 0.884 | 0.714 | 0.11 |
| LYSIMNIA | 3 | 3 | 0.71 | 0.906 | 0.621 | 0.094 |
| MAMERCUS_largo | 8 | 28 | 0.761 | 0.881 | 0.759 | 0.054 |
| NOTABILIS | 2 | 1 | 0.182 | 0.968 | 0.421 | 0.004** |
| ORESTES_largo | 6 | 15 | 0.61 | 0.888 | 0.736 | 0.006** |
| PAVANA | 3 | 3 | 0.971 | 0.909 | 0.649 | 0.733 |
| POSTMAN_superlargo | 7 | 21 | 0.886 | 0.886 | 0.731 | 0.5 |
| RICINI_largo | 23 | 253 | 0.781 | 0.876 | 0.818 | 0.011* |
| VANILLAE_largo | 13 | 78 | 0.796 | 0.874 | 0.791 | 0.06 |
| WALLACEI | 9 | 36 | 0.67 | 0.884 | 0.775 | 0.005** |
| XENOCLEA | 2 | 1 | 0.395 | 0.966 | 0.452 | 0.035* |
| **Inter-tribe groups** | | | | | | |
| DERCYLLIDAS | 2 | 1 | 0.261 | 0.999 | 0.653 | 0.002** |
| ELZUNIA_largo | 7 | 10 | 0.664 | 0.951 | 0.839 | 0.001*** |
| EURIMEDIA | 36 | 35 | 0.979 | 0.94 | 0.874 | 0.826 |
| EXCELSA | 22 | 72 | 0.805 | 0.94 | 0.896 | 0.001*** |
| LYSIMNIA | 8 | 15 | 0.728 | 0.944 | 0.864 | 0.001*** |
| MAMERCUS_largo | 91 | 664 | 0.861 | 0.939 | 0.916 | 0.001*** |
| ORESTES_largo | 30 | 144 | 0.748 | 0.939 | 0.905 | 0.001*** |

Beyond spatial congruence, patterns of niche convergence in Heliconiine butterflies were supported indifferently with the ‘split’ classification (**Fig S7.a**; Permutation test: $MCD_{obs}$ = 0.684, $MCD_{null}$ Q5% = 0.882, $p \leq 0.001$), or the ‘lumped’ classification (**Fig S10.a**; Permutation test: $MCD_{obs}$ = 0.866, $MCD_{null}$ Q5% = 0.869, $p = 0.045$). Similarly, we found a signal for niche convergence between pairs of Heliconiini and Ithomiini taxa (i.e., inter-tribes) sharing similar phenotypes for both the ‘split’ classification (**Fig S7.c**; Permutation test: $MCD_{obs}$ = 0.725, $MCD_{null}$ Q5% = 0.866, $p \leq 0.001$) and the ‘lumped’ classification (**Fig S10.b**; Permutation test: $MCD_{obs}$ = 0.681, $MCD_{null}$ Q5% = 0.841, $p \leq 0.001$). At group-level, we detected a significant signal for spatial congruence in 4 out of 15 (26.6%) 'lumped' phenotypic groups of heliconiines (**Table S6**). This proportion rises to 3 out of 7 (42.8%) 'lumped' groups for inter-tribe phenotypic groups shared between the heliconiine and ithomiine butterflies (**Table S6**).

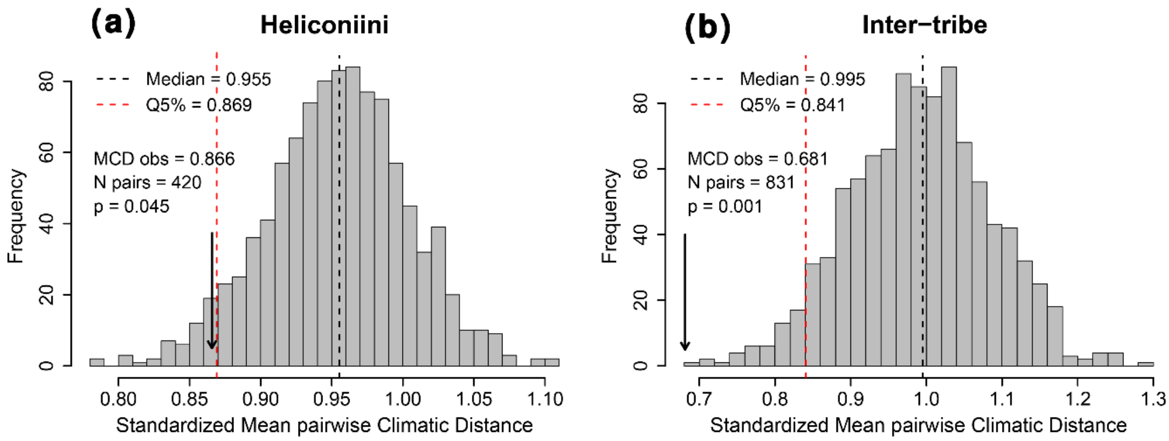


**Figure S10: Null distributions for niche convergence tests within phenotypic groups using the 'lumped' classification. (a)** Heliconiini **(b)** Ithomiini adapted from Doré et al. (1) **(c)** Inter-tribe phenotypic groups. Phylogenetic comparative tests with 1000 simulations of neutral niche evolution (Brownian Motion model with Pagel's lambda = 0.798). Standardized Mean pairwise Climatic Distances (MCD) quantify the degree of dissimilarity in niche centroids between pairs of OMUs. An observed mean MCD value lower than the 5% quantile (Q5%) of the null distribution supports the overall niche convergence of phenotypically similar species.

**Table S6: Tests for niche convergence within each phenotypic group using the 'lumped' classification.** Standardized Mean pairwise Climatic Distances (MCD) quantify the degree of dissimilarity in niche centroids between pairs of OMUs. An observed mean MCD value lower than the 5% quantile (Q5%) of the null distribution obtained through simulation of the neutral evolution of the niche supports the niche convergence of phenotypically similar species within a phenotypic group. Tests for inter-tribe phenotypic groups are carried out between pairs of heliconiine vs. ithomiine OMUs. $MCD_{obs}$ = Observed MCD. $MCD_{Q50}$ = Median of the null distribution. $MCD_{Q5}$ = Quantile 5% of the null distribution = statistical significance threshold for α = 0.05. P-value for the spatial congruence test is based on the rank of the $MCD_{obs}$ among the null distribution obtained through neutral evolution of the niche (N = 1000): * = p < 0.05; ** = p < 0.01; *** = p ≤ 0.001.

| PHENOTYPIC GROUP | N units | N pairs | $MCD_{obs}$ | $MCD_{Q50}$ | $MCD_{Q5}$ | p-value |
|---|---|---|---|---|---|---|
| **Heliconiini groups** | | | | | | |
| CHARITHONIA | 1 | 0 | | | | |
| CHESTERTONII | 1 | 0 | | | | |
| CYDNIDES_largo | 7 | 21 | 0.532 | 0.899 | 0.605 | 0.02* |
| DIDO | 5 | 10 | 0.767 | 0.774 | 0.457 | 0.484 |
| ELZUNIA_largo | 5 | 10 | 1.531 | 0.888 | 0.541 | 0.996 |
| EURIMEDIA | 1 | 0 | | | | |
| EXCELSA | 4 | 6 | 1.877 | 1.077 | 0.593 | 0.989 |

| | | | | | | |
|---|---|---|---|---|---|---|
| HEURIPPA | 1 | 0 | | | | |
| LEUCE_largo | 5 | 10 | 1.288 | 1.005 | 0.603 | 0.859 |
| LYSIMNIA | 3 | 3 | 0.36 | 0.943 | 0.469 | 0.016* |
| MAMERCUS_largo | 8 | 28 | 0.822 | 0.972 | 0.675 | 0.204 |
| NOTABILIS | 2 | 1 | 0.495 | 1.212 | 0.315 | 0.099 |
| ORESTES_largo | 6 | 15 | 0.452 | 0.952 | 0.626 | 0.003** |
| PAVANA | 3 | 3 | 0.633 | 0.821 | 0.377 | 0.267 |
| POSTMAN_superlargo | 7 | 21 | 1.106 | 1.03 | 0.695 | 0.626 |
| RICINI_largo | 21 | 210 | 0.851 | 0.952 | 0.785 | 0.153 |
| VANILLAE_largo | 10 | 45 | 0.967 | 0.978 | 0.699 | 0.484 |
| WALLACEI | 9 | 36 | 0.737 | 0.882 | 0.622 | 0.18 |
| XENOCLEA | 2 | 1 | 0.056 | 1.221 | 0.313 | 0.001*** |
| **Inter-tribe groups** | | | | | | |
| ELZUNIA_largo | 7 | 10 | 1.061 | 0.939 | 0.535 | 0.673 |
| EURIMEDIA | 34 | 33 | 0.675 | 0.94 | 0.618 | 0.099 |
| EXCELSA | 21 | 68 | 1.03 | 1.003 | 0.786 | 0.565 |
| LYSIMNIA | 7 | 12 | 0.339 | 0.97 | 0.653 | 0.002** |
| MAMERCUS_largo | 80 | 576 | 0.689 | 0.991 | 0.826 | 0.002** |
| ORESTES_largo | 28 | 132 | 0.469 | 0.997 | 0.764 | 0.001*** |
| ELZUNIA_largo | 7 | 10 | 1.061 | 0.939 | 0.535 | 0.673 |

### Appendix 8: ODMAP

#### Overview

##### Authorship

**Authors:** Eddie Pérochon, Neil Rosser, Krzysztof Kozak, W. Owen McMillan, Blanca Huertas, James Mallet, Jonathan Ready, Keith Willmott, Marianne Elias, Maël Doré.

**Contact:** Maël Doré ;

**Title:** Müllerian mimicry in Neotropical butterflies: One mimicry ring to bring them all, and in the jungle bind them

##### Model objective

**Model objectives:** Mapping and interpolation. We mapped current potential distribution of subspecies of heliconiine butterflies.

**Target output:** Meeting our objectives required several steps in the post-processing of model outputs. (i) We obtained environmental suitability maps depicting potential distributions from SDM for each subspecies. (ii) We aggregated predictions to derive distributions for Operational Mimicry Units (OMUs; 4), species and mimetic groups as likelihood of presence of at least one subspecies from the OMU/species/mimetic group. (iii) We obtained richness maps as stacked-SDMs from species and mimetic group maps. (iv) We computed various taxonomic, phylogenetic and phenotypic diversity and rarity indices from the previous richness maps.

##### Focal Taxon

**Focal Taxon:** Our study group was the longwing butterflies in the tribe Heliconiini Swainson, 1822 (Nymphalidae: Heliconiinae). This clade contains ca. 8 genera, 77 species and 457 subspecies (8, 9), but see Núñez et al. (10) for recent proposed taxonomic splits). Our study includes the 439 subspecies (96.1 %) with available georeferenced occurrences.

##### Location

**Location:** Americas, from Argentina to Canada, including the Caribbean region, encompassing the whole range of the Heliconiini tribe. Most diversity falls within the Neotropics with only a few lineages venturing in the Nearctic region.

##### Scale of Analysis

**Spatial extent:** Longitude 130° E - 30° E, Latitude 38° S – 50° N

**Spatial resolution:**

Community boundaries were defined as grid cell of 0.25° x 0.25° = 27.8km x 27.8km.

**Temporal extent:** Field surveys were conducted in the last decades. The dataset is complemented with historical records that span from the 19th century to present, with the majority collected within the last 30 years.

**Temporal resolution:** We modeled distributions under current environmental conditions: we retrieved bioclimatic data for the 2000's decade, and forest cover data for the year 2010.

##### Biodiversity data

**Observation type:** Georeferenced occurrences from field surveys and data from specimens in private and museum collections

**Response data type:** A curated set of 67,563 subspecies-locality records as presence data were screened to yield 18,841 subspecies-grid-cell records after removing duplicate records from single grid cells. We drew pseudo-absences from those occurrences in a target group strategy. See details in Data.

##### Predictors

**Predictor types:** bioclimatic, topographic (elevation), and habitat (forest cover)

##### Hypotheses

**Hypotheses:**

Heliconiini species inhabit either forests or savannah across Americas (8). Thus, their distribution is expected to be widely influenced by the local availability of forest cover. Likewise, elevation has been shown to shape the broadscale patterns of Heliconiini diversity (11). We also used climatic layers as predictors in an exploratory way because climate is known to be an important driver of species distributions at a continental scale in general (12).

##### Assumptions

**Model assumptions:** We assumed that (i) relevant ecological drivers (or proxies) of species distributions are included, (ii) detectability does not change across environmental gradients, (iii) predictor measurements are free of error, (iv) the species are at equilibrium with their environment, (v) sampling is sufficient and representative, and (vi) environmental suitability outputs are valuable proxies to estimate potential distributions.

##### Algorithms

**Modelling techniques:** We employed three different machine learning algorithms: Random Forest (RF), Generalized Boosted Models (GBM) also known as Boosted Regression Trees (BRT), and Artificial Neural Networks (ANN).

**Model complexity:** We kept model settings to the default settings in *biomod2* v.3.4.6, keeping a balance between flexibility of the response curves and overfitting (13). The only exceptions were the minimum size of leaves that was lowered to two instead of five, and the fraction of observations used at each step (0.7 instead of 0.5) to allow tentative runs in GBM for OMUs with low sample size.

**Model ensembles:** We stacked all models meeting our validation thresholds to produce a single "ensemble" model per subspecies. We computed the ensemble as the median rather than the mean to limit the influence of models with extreme outputs. We did not use a weighting scheme since we considered our evaluation metric (i.e., Jaccard index) a suitable metric to discard low quality models, but not adequate to rank best models in the context of pseudo-absences data (14).

#### Workflow

**Model workflow:** We fitted SDMs for 364 subspecies (91.4 %) representing 75 species (97.4%), for which we had at least six occurrences available. We included the remaining 75 subspecies in stacks as binary raster of presence-absences. Workflow also differed between the 196 subspecies with restricted sample size (6 ≤ N < 30) and the 168 subspecies with large sample size (N ≥ 30). For restricted sample size, we kept all occurrences for calibration and validation and draw 10 independent pseudo-absences sets. For large sample sizes, we drew three independent pseudo-absences sets combined with 3-fold spatial block cross-validation to assess predictive model performance. We selected valid models for ensemble based on maximized Jaccard indices and plausibility checks. Ensemble predictions were derived using ensemble medians. We clipped final outputs with subspecies-specific buffered alpha-hulls and Andean region masks to constrain the extent of possible distributions to reasonable areas. We derived OMU, species and mimetic group maps from the subspecies maps as the likelihood to find at least one of the related subspecies in the community. Final post-processing step consisted in the computation of six diversity and rarity indices based directly on the species or mimetic group maps. This workflow is depicted in as a chart adapted from Doré et al. (4) in **Figure S11**.

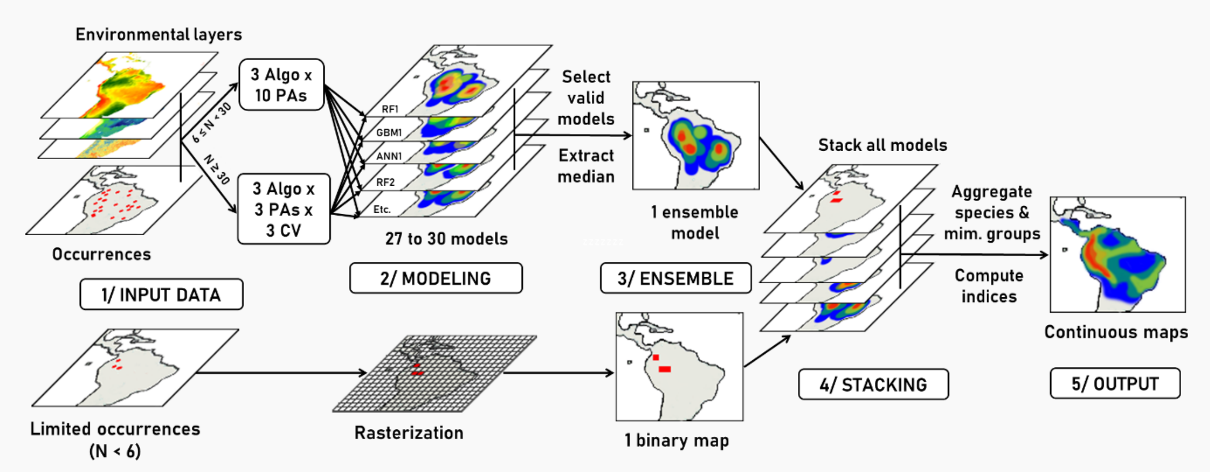


**Figure S11: SDM workflow chart depicting the different steps of the analysis performed in this study.** Depending on sample size, modeling steps and settings differed. Clipping step to constrain SDM projections to plausible distribution ranges is not shown on the chart. Algo = algorithms used in the study, namely random forest (RF), gradient boosting models (GBM), and artificial neural networks (ANN); PAs = pseudo-absences sets; CV = cross-validation folds. Modified from ODMAP in Doré et al. (4).

#### Software

**Software:** R version 3.6.2 (15) with packages *raster* 3.0-12 (16), *biomod2* 3.4.6 (17), *sf* 0.9-0 (18), *blockCV* 2.1.1 (19), *alphahull* 2.2 (20), and others.

**Code availability:** All scripts are provided on GitHub at https://github.com/EddiePerochon/Heliconiini_Diversity

**Data availability:** Occurrence data and mimetic group delimitation used for modeling are available from Zenodo at 10.5281/zenodo.10906853 and 10.5281/zenodo.10903197. All subspecies/OMU/species/mimetic group distribution maps are available from Zenodo at 10.5281/zenodo.10903661.

### Data

##### Biodiversity data

**Taxon names:** Heliconiini tribe. All subspecies are listed in the phenotypic classification available in 10.5281/zenodo.10903197.

**Taxonomic reference system:** Current names in use as listed in Jiggins & Lamas (8).

**Ecological level:** Subspecies, and species in case of taxa with a unique or no subspecies described.

**Data sources:** Dataset of georeferenced occurrences is a compilation of fieldwork data from N. Rosser (Harvard University, USA), J. Mallet (Harvard University, UK), K. Kozak (STRI, Panama) and O. McMillan (STRI, Panamá) obtained over the past decades.

Additionally, the dataset comprises records from museums and private collections compiled by N. Rosser. Data sources are summarized in Table 1 in Rosser et al. (11). The main contributors (>3000 records each) are the Florida Museum of Natural History, Gainesville (FLMNH), the Natural History Museum, London (NHMUK), the Tropical Andean Butterfly Diversity Project, CONABIO, Mexico, and the Museo de Historia Natural, Universidad Nacional Mayor de San Marcos, Lima (MUSM).

The dataset of the curated 67,563 subspecies-locality records used for distribution modeling is available from Zenodo at 10.5281/zenodo.10906853.

**Sample size:** Sample size for each subspecies, after spatial filtering, varied widely from 1 to 565. We employed a different modeling scheme for subspecies falling into different sample size categories. We classified the 75 subspecies (17.0%) with sample size lower than six as “rasterized” and did not go through the SDM process. We labelled as “restricted” the 196 subspecies (44.6%) with sample size between 6 and 29. We modeled these “restricted” subspecies without cross-validation. The 168 subspecies (38.3%) with sample size greater or equal to 30 underwent the full SDM process. See workflow in **Fig. S11** above.

**Scaling:** We removed spatial duplicates by applying spatial filtering on a 0.25° x 0.25° grid used as final modeling resolution whose pixels defined our virtual communities. As a consequence we obtained 18,841 subspecies-grid-cell records.

**Cleaning:** We examined the presence of geographic and environmental outliers prior to modeling. We considered as geographic outliers all occurrences of a specific subspecies with no other neighboring points in a 1000km buffer area. These outliers were further scrutinized to decide case by case to retain or discard those points from the dataset if considered erroneous or not.

We automatically removed occurrences with significant Mahalanobis distance (21) from other points in the environmental space. Those points could be either errors, or real abnormal occurrences, caused by temporary migration of individuals following an extreme climatic event (e.g., individuals migrating temporarily up mountain slopes following an extreme heat event).

In any case, those occurrences cannot be considered helpful to model the global species distribution based on the local average climate and were therefore discarded.

**Pseudo-absence data:** We generated pseudo-absences using a target-group strategy (22), employing sampling sites where other subspecies have been detected but not the targeted subspecies as a pool for drawing pseudo-absences. In doing so, we increased the likelihood for the targeted subspecies to be effectively not present in our pseudo-absence sites, a critical aspect in order to produce quantities that approach the actual probability of occurrence of the entity modeled as output, as we intended to do (23). This approach also allowed us to use confidently the Jaccard index as an evaluation metric to discard poorly performing models from the ensemble models despite this measurement being based on confusion matrix, thus designed primarily for presence-absence data.

Additionally, in order to minimize even more the risk of assigning wrongly a pseudo-absence in an actual occupied site we applied a minimum buffer of 1° (111.32 km at the equator) to discard all sites within this minimum range of a presence point from the potential pseudo-absence pool. Finally, to prevent selecting pseudo-absences too far from any presence points while avoiding having to decide a global arbitrary maximum threshold, we weighted the probability for sites to be selected by their inverse distance to any presence point. Therefore, we ensure our pseudo-absences were likely to represent real absences, while at the same time avoiding to extensively sample too far beyond the range of a species, where absences are likely to occur because of non-bioclimatic reasons (e.g., 24).

Following recommendations from Barbet-Massin et al. (25) for machine-learning algorithms, we drew a number of new pseudo-absences equal to the number of presences recorded for the target subspecies, for each run. For each subspecies with a restricted sample size ($6 \leq N < 30$), we ran ten independent replicates for each algorithm leading to a total of 30 models per subspecies. For each subspecies with a large enough sample size ($N \geq 30$), we ran three independent replicates for each algorithm leading to a total of 27 models per subspecies (i.e., 3 algorithms * 3 pseudo-absence sets * 3 CV-folds), once the 3-fold spatial blocks CV was applied.

#### Data partitioning

**Validation data:** We split data between training set and validation sets only for subspecies with sample size ≥ 30.

For models with limited sample size ($6 \leq N < 30$), we decided to keep all data points in our calibration set in order to yield better estimates from SDMs with low sample size. Using a partition scheme would have left fewer points for calibration, decreasing the already scarce information available to yield proper SDMs, and even fewer for validation which would have become meaningless (26). Thus, we evaluated model performances with the same dataset used for calibration (« resubstitution » in Roberts et al. (27)). To compensate for non-independence between our calibration sets and validation sets, we used conservative high thresholds to select models with valid performance to keep for the final ensemble.

For models with sufficient sample size (N >= 30), we applied a 3-folds cross-validation (CV) strategy with spatial blocks to define our calibration and validation sets. Spatial blocks CV allows to partition dataset into spatially independent blocks that ensure the predictive error of the model is not underestimated because of spatial autocorrelation between calibration and validation sets (27). We defined our folds for each subspecies dataset of presences combined with each independent draw of pseudo-absences using the *spatialBlock* function in the R package *blockCV* 2.1.1 (19).

**Test data:** No truly independent dataset was available.

#### Predictor variables

**Predictor variables:**

Climate is known to widely influence large-scale patterns of species distribution (28). We selected as predictors four bioclimatic variables among the 19 available in the WorldClim 2.1 online database (29): mean annual temperature, mean diurnal range, total annual precipitation, and precipitation seasonality. We selected this subset of climate predictors following two aims: (1) representativity of the whole variance in our study region, and (2) ease of ecological interpretations. Each variable was included in a different group of intercorrelated variables when performing hierarchical clustering. Within each group, we selected the most adequate variables for ease of ecological interpretation of effects of climate on species distribution.

We used elevation as it has been shown to shape the broadscale patterns of Heliconiini diversity (11).

We used percentage of forest cover as our habitat/land use type predictor since Heliconiini species are known to be found either in forests or savannah across Americas (8).

**Data sources:**

Bioclimatic predictors were obtained from WorldClim v2.1 (29) for the latest period available at the time of our modeling process (1970-2000).

We retrieved elevation from the SRTM Dataset v.4.1 (30; http://srtm.csi.cgiar.org/, accessed on 03/26/2019).

We extracted the percentage of land cover per pixel from the Landsat Tree Cover Continuous Fields dataset (31) for the year 2010, accessible through Google Earth Engine (GLCF: Landsat Tree Cover Continuous Fields in the Earth Engine Data Catalog, accessed on 03/26/2019). The GLCF tree cover layers contain estimates of the percentage of horizontal ground covered by woody vegetation.

**Spatial extent:** We clipped all rasters to our study area: Longitude 130° E - 30° E, Latitude 38° S – 50° N.

**Spatial resolution:** We obtained GLCF and WorldClim data at a resolution of 5min of arc, while SRTM had a resolution of 90m. We aggregated all predictor variables to our final model resolution (0.25°) prior to modeling.

**Coordinate reference system:** Data were retrieved under WGS84 (EPSG:4326). All rasters were projected to Mollweide projections (ESRI:54009) priori to modeling in order to ensure grid cells represented similar areas.

**Temporal extent:**

We downloaded WorldClim data averaged for the 1970-2000 decades, and GLCF for the year 2010.

**Data processing:** We aggregated all predictor variables to our final model resolution (0.25°) prior to modeling. We harmonized final predictor rasters to display missing data in pixels where at least one predictor was lacking information to avoid modeling points with partial environmental information.

**Dimension reduction:** We selected four bioclimatic variables among the 19 bioclimatic variables available in WorldClim in order to reduce multicollinearity among bioclimatic predictors, and limit the complexity of the models to a reasonable number of predictors (six in total).

We selected bioclimatic variables as the results of a hierarchical agglomerative clustering on Spearman's rho correlation coefficients, using a complete linkage method with the function *hclust* in R base package. We applied a cutoff of $|\rho| > 0.7$ (32, 33) on the resulting dendrogram to highlight groups of multicorrelated variables. Then, we selected only one variable in each group. Selection criteria for retaining variables were (1) their ease to interpret as an ecological factor (e.g., "mean temperature" rather than the "mean temperature of the driest quarter") and (2) their high correlation with the axis of a global PCA run on all 19 variables. The final four variables included in the models were mean annual temperature, mean diurnal range, total annual precipitation, and precipitation seasonality.

##### Transfer data

We interpolated our final maps of environmental suitability depicting potential distributions with the same environmental rasters as the one used for modeling. Thus, the transfer data is the same as the predictors.

#### Model

##### Variable pre-selection

See Dimension reduction in Data section.

##### Multicollinearity

See Dimension reduction in Data section.

##### Model settings

**Model complexity:** We kept model settings to the default settings in *biomod2* v.3.4.6, since models with intermediate levels of complexity have been shown to perform best (33), keeping a balance between flexibility of the response curves and overfitting (13). The only exceptions to default settings were the minimum size of leaves that was lowered to two instead of five, and the fraction of observations used at each step (0.7 instead of 0.5) to allow tentative runs in GBM for subspecies with low sample size.

**BRT/GBM settings:** distribution (bernoulli), nTrees (2500), interactionDepth (7), shrinkage (0.001), bagFraction (0.7), trainFraction (1), n.minobsinnode (2), CV.folds (3)

**randomForest settings:** ntree (500), mtry (2), maxnodes (n.obs), sampsize (n.obs), replace (TRUE)

**ANN settings:** nbCV (5), maxit (200)
size = 2, 4, 6, or 8. Optimized by CV for best AUC.
decay = 0.001, 0.01, 0.05, or 0.1. Optimized by CV for best AUC.

**Model extrapolation:** Extrapolation was possible but remained limited since we constrained final outputs inside the buffer around known presence points.

##### Model estimates

**Variable importance:** We assessed variable importance for each calibrated model within the R package *biomod2* 3.4.6 (17) by looking at the correlation between predictions obtained from the real data and predictions from data with randomized values for each variable evaluated.

##### Model selection - model averaging - ensembles

**Model selection:** We discarded all models that did not reach our thresholds for model quality prior to ensemble (See Performance statistics in Assessment section for details on evaluation metric choice). We set our threshold to a minimum Jaccard index of 0.6 for complete models (N >= 30), and 0.95 for restricted models ($6 \leq N < 30$). The threshold for subspecies with "restricted" sample size was more conservative since they were evaluated on the calibration set, while complete models were evaluated on spatially independent validation sets. We chose those thresholds since they ensured each subspecies retained at least 5 models for the ensemble, while keeping quality standard to a high value. We conducted additional plausibility checks by inspecting the response curves of each variable for each model following an automatic procedure, completed with manual checks (See plausibility checks in Assessment for details). We discarded from the ensemble models holding at least one response curve with a non-ecologically plausible shape.

**Model ensembles:** We stacked all models meeting our validation thresholds to produce a single "ensemble" model per subspecies. We computed the ensemble as the median rather than the mean to limit the influence of models with extreme outputs. We did not use a weighting scheme since we considered our evaluation metric (i.e., Jaccard index) a suitable metric to discard low quality models, but not adequate to rank best models in the context of pseudo-absence data (14).

##### Analysis and Correction of non-independence

**Spatial autocorrelation:** We applied spatial blocks CV to account for spatial autocorrelation among calibration and validation sets for models with sufficient sample size (N >= 30).

##### Threshold selection

**Threshold selection:** We did not apply a threshold on the final continuous outputs prior stacking since it has been proven that thresholding could introduce bias leading to overestimation of species richness (34). For instance, we simply estimated species richness as the sum of species environmental suitability maps depicting potential distributions as proxies of occurrence probabilities.

#### Assessment

##### Performance statistics

**Metric choice:** In order to evaluate model performance, we chose to use the Jaccard index, an ecological index of similarity which can be directly interpreted as the spatial overlap between the observed distribution (valid predicted presences as true positives (TP), and missed presences as false negatives (FN)) and predicted distribution (valid predicted presences as true positives (TP), and erroneous predicted presences as false positives (FP)). Thus, for each model we retained the maximum Jaccard index obtained for a model specific optimized threshold, and computed as TP/(TP + FN + FP). Contrary to the TSS, the Jaccard index prevents overestimation of model performance caused by the inflation of true negatives based on pseudo-absences drawn far from presences, and appeared to be not biased by prevalence (14).

Additionally, despite being primarily designed for presence-absence data, we used the Jaccard index as evaluation metric because: (1) we did not have enough occurrence data to use presence-only evaluation metrics such as the Boyce index (35) for most of our subspecies; (2) we carefully selected our pseudo-absences in a target-group strategy (22) to maximize the probability for our pseudo-absences to be real absences, thus we are confident the Jaccard index remains informative to discard poorly performing models.

**Performance on training data:** We set to a high 0.95 the threshold to meet required quality for "restricted" models evaluated directly on the calibration set because of low sample size.

**Performance on validation data:** We set to 0.6 the threshold to meet required quality for "complete" models (N >= 30) evaluated using spatial CV-blocks.

Following our criteria, we retained 87.3% of sub-models run with Random Forest algorithms, 96.0% of Gradient Boosted Models, and 56.7% of Artificial Neural Networks (**Fig. S12**). All ensemble models retained at least five sub-models, allowing to provide final predictions based on ensemble models accounting for uncertainties associated with modeling choices.

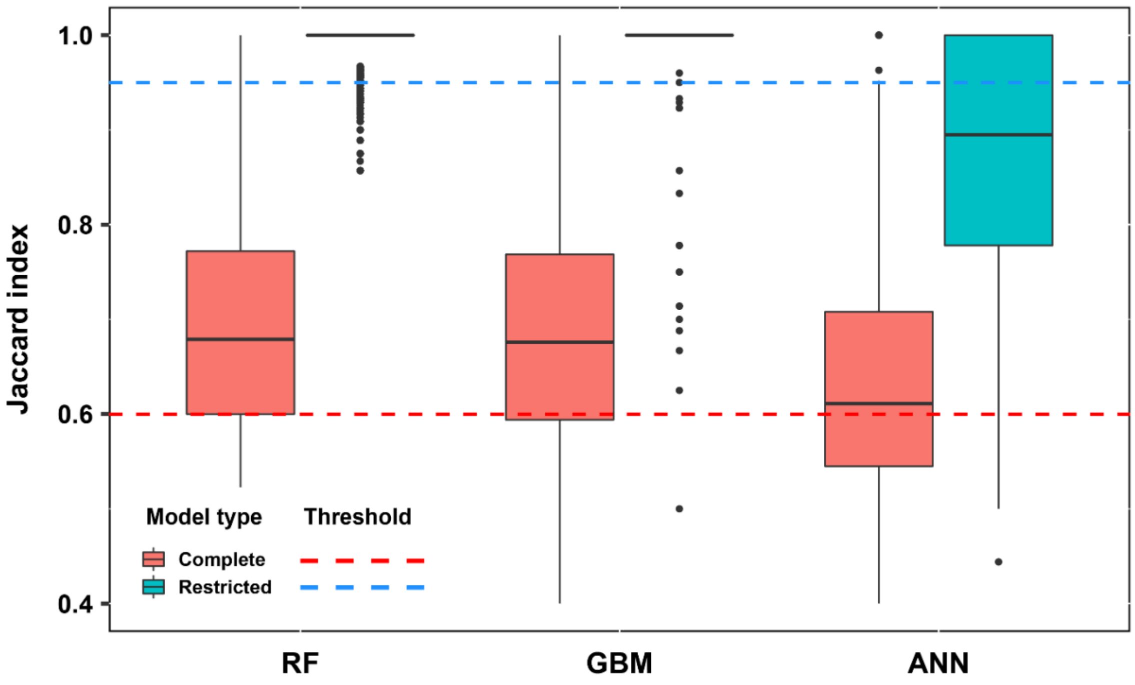


**Figure S12: Evaluation of sub-model performances based on Jaccard indices.** Sub-models are grouped by type of algorithms: Random Forest (RF), Gradient Boosted Models (GBM), and Artificial Neural Networks (ANN). Two distinct thresholds (dashed lines) were used to discard from ensemble models the sub-models with insufficient performance depending on the type of evaluation set used: 0.95 when evaluated on the training set data ("restricted models", in blue), 0.6 when evaluated on a validation set designed from spatial CV-blocks ("complete models", in red).

##### Plausibility check

**Response shapes:** We designed an automatic procedure to check for multimodality and positive quadratic relationships in the response curve of all variables for all models since such relationships would have low ecological plausibility. We assessed multimodality through Hartigan's dip test using the R package *diptest* 0.75-7 (36). We inspected case by case the response curves highlighted by the automatic procedure and then manually removed models holding non-plausible response curves based on expert judgement.

#### Prediction

##### Prediction output

**Prediction unit:** Models produced environmental suitability maps depicting potential distribution for each subspecies.

**Clipping:** We clipped all final ensemble model per unit using a subspecies-specific buffer.

We clipped all final ensemble models to constrain the extent of possible distribution of each subspecies to a reasonable area accordingly to the limited migration abilities of our butterflies, and the degree of certainty we had about the range of the species based on the spread of occurrence points. Thus, we computed alpha-hulls (α = 1000 km) using the function *ahull* from the R package *alpha-hull* 2.2 (20) encompassing all occurrence points for each subspecies in order to design a smooth surface able to engulf occurrences points but that could also generate automatically disjointed distributions when needed. We choose an alpha parameter of 1000 km (diameter of the circles used to draw the alpha-hull) to be coherent with our threshold for detection of outliers. In parallel, we computed the 80% quantile for the distance to the closest occurrence points among occurrence points of each subspecies. This measure is to be seen as a measurement of how confident we are that our records cover extensively the range of the subspecies studied. The rationale is that a subspecies with a clustered set of occurrences is more likely to describe accurately the global range of this subspecies, while a subspecies with a more dispersed set of occurrence points could signal a lack of information, or a subspecies with a wide range. Therefore, we added to our alpha-hull a buffer corresponding to the max value between this subspecies-specific parameter and a distance of 1° assumed to represent a conservative limit for Heliconiini dispersion abilities. We used these final polygons to restrict our SDMs predictions for each subspecies.

In the specific case of the Andean region, strong environmental gradients can be found following the slopes, leading to potentially suitable areas on both sides of the Cordilleras across limited distances. To avoid false predictions of subspecies known with reasonable confidence to be restricted to one side of the Cordilleras, we cropped the final maps applying a set of two polygons corresponding respectively to each side of the Andean mountain ranges. We built those polygons by aggregating watersheds retrieved from a Digital Elevation Model in ArcGIS. In practice, if a subspecies presented occurrences falling only in one of the two polygons, we used the other to crop out the final map of this subspecies.

**Aggregating to higher level:** Exploiting the 439 environmental suitability maps depicting potential distributions for each subspecies, we built potential distribution maps at OMU, species and mimetic group levels.

We assumed outputs from SDMs relate to likelihood of presence of each subspecies. Then, for each community, the likelihood of presence of an OMU, a species, or a mimetic group was computed as the likelihood to find at least one of the related subspecies in the community such as

$$p_{o/s/m} = 1 - \prod(1 - p_i) \qquad (1)$$

where $p_{o/s/m}$ is the likelihood of presence of the OMU, species or mimetic group, and $p_i$ the likelihood of presence of each subspecies of this OMU, species or mimetic group.

**Index computation:** A final post-processing step consisted in the computation of six diversity and rarity indices based directly on the species or mimetic group maps. For all computation, we used the continuous outputs from SDM since binarization to presence-absences usually degrades inference and can introduces bias in community richness evaluation (23, 34).

##### Uncertainty quantification

**Algorithmic uncertainty:** We accounted for algorithmic uncertainty by applying an ensemble approach averaging over three different SDM algorithms.

**Input data uncertainty:** We accounted for uncertainty in input data by applying an ensemble approach averaging over three (for subspecies with additional spatial blocks CV) or ten (for subspecies without CV) different pseudo-absences draws.

**Novel environments:** Predictions to novel environments were limited since we interpolate maps inside a buffer encompassing known presence points.
